# Pinhead antagonizes Admp to promote notochord formation

**DOI:** 10.1101/2021.04.21.440817

**Authors:** Keiji Itoh, Olga Ossipova, Sergei Y. Sokol

**Affiliations:** Department of Cell, Developmental and Regenerative Biology, Icahn School of Medicine at Mount Sinai, New York

**Keywords:** proteomics screen, BMP4, Admp, Smad1 phosphorylation, Pnhd, gastrulation, secretome, notochord, Chordin, Xenopus.

## Abstract

Dorsoventral patterning of a vertebrate embryo critically depends on the activity of Smad1 that mediates signaling by several BMP proteins, anti-dorsalizing morphogenetic protein (Admp), and their antagonists. Pinhead (Pnhd), a cystine-knot-containing secreted protein, is expressed in the ventrolateral marginal zone during *Xenopus* gastrulation, however, its molecular targets and signaling mechanisms have not been fully elucidated. An unbiased mass spectrometry-based screen of the gastrula *secretome* identified Admp as a primary Pnhd-associated protein. We show that Pnhd binds Admp and inhibits its ventralizing activity by reducing Smad1 phosphorylation and suppressing its transcriptional targets. By contrast, Pnhd did not affect the signaling activity of BMP4. Importantly, the Admp gain-of-function phenotype and phospho-Smad1 levels have been enhanced after Pnhd depletion. Furthermore, Pnhd strongly synergized with Chordin and a truncated BMP4 receptor in the induction of notochord markers in ectoderm cells, and Pnhd-depleted embryos displayed notochord defects. Our findings suggest that Pnhd binds and inactivates Admp to promote notochord development. We propose that the interaction between Admp and Pnhd refines Smad1 activity gradients during vertebrate gastrulation.

## Introduction

The specification of the dorsoventral axis and the three germ layers in vertebrate embryos is controlled by several signaling pathways triggered by Nodal, FGF and Wnt ligands (De Robertis and Kuroda, 2004; Harland and Gerhart, 1997; Kiecker et al., 2016). During gastrulation, the activity of the Spemann organizer results in the induction of the neural tissue in the ectoderm, whereas the mesoderm becomes subdivided into the notochord, paraxial (somites), intermediate (lateral plate, kidney) and ventral (blood) mesoderm (Kimelman, 2006; Niehrs, 2004). Both neuralization of the ectoderm and dorsalization of the mesendoderm primarily rely on a Smad1 activity gradient established by ventrolaterally expressed <u>b</u>one <u>m</u>orphogenetic <u>p</u>roteins (BMP4 and BMP7 ligands) and organizer-specific BMP antagonists Chordin, Noggin and Follistatin (De Robertis and Kuroda, 2004; Harland and Gerhart, 1997; Tuazon and Mullins, 2015).

Genetic and molecular studies implicated BMPs and their antagonists in the regulation of dorsoventral patterning during vertebrate gastrulation. BMPs specify ventral fates in a dose-dependent manner by stimulating the phosphorylation of Smad1/5/8 (referred to as Smad1, for simplicity). By contrast, Chordin secreted by the organizer, binds and inhibits BMPs to promote dorsal fates. Both Chordin and BMPs diffuse in the extracellular space and form opposing activity gradients in the embryo (Dale et al., 1992; De Robertis and Kuroda, 2004; Piccolo et al., 1996). Phospho-Smad1 is distributed along the dorsoventral axis in a graded fashion that corresponds to the putative BMP activity gradient (Faure et al., 2000; Schohl and Fagotto, 2002). In amphibians, <u>a</u>nti-<u>d</u>orsalizing <u>m</u>orphogenetic <u>p</u>rotein (Admp), a dorsally expressed BMP ligand, also stimulates Smad1 phosphorylation to expand the zygotic BMP gradient (Dosch and Niehrs, 2000; Joubin and Stern, 1999; Moos et al., 1995; Reversade and De Robertis, 2005; Willot et al., 2002). Thus, the balance between the activators and inhibitors of Smad1 signaling is essential for dorsoventral patterning.

Pinhead is a conserved zygotic secreted protein implicated in *Xenopus* embryonic axis development (Kenwrick et al., 2004). Pnhd contains conserved cystine knot motifs and is induced in the ventrolateral marginal zone by Wnt and FGF signaling during gastrulation (Kjolby and Harland, 2017; Ossipova et al., 2020). Pnhd has been proposed to function synergistically with FGF, Nodal and BMP pathways during mesoderm development (Ossipova et al., 2020; Yan et al., 2019). Nevertheless, Pnhd-interacting proteins and signaling mechanisms remain to be characterized in further detail.

In this study, we initiated an unbiased mass spectrometry screen for *Xenopus* proteins that associate with Pnhd in the extracellular space. We identified the top candidate protein as Admp, an organizer-specific agonist of the BMP pathway. We find that Pnhd binds to Admp and inhibits its ventralizing activity mediated by phospho-Smad1 signaling. Moreover, Pnhd together with Chordin induces dorsal mesoderm markers. We also find that Pnhd is required for *Xenopus* notochord development consistent with the idea that Pnhd functions as Admp antagonist. We propose that the interaction of Pnhd and Admp in the marginal zone buffers Smad1 activity gradient during dorsoventral patterning.

## Results

### Screening for Pnhd-interacting proteins in the gastrula secretome

Many cystine knot proteins, such as Cerberus or Gremlin (Hsu et al., 1998; Piccolo et al., 1999; Sasai et al., 1994), associate with other secreted proteins rather than cell surface receptors. Since Pnhd is readily secreted by cell lines and dissociated *Xenopus* gastrula cells (Ossipova et al., 2020; Yan et al., 2019), we decided to perform a screen for Pnhd-interacting partners in the extracellular space. This approach allows to enrich putative Pnhd-binding molecules by eliminating the abundant yolk proteins and common cytoplasmic contaminants, such as actin or tubulin.

In our gastrula secretome screen, the conditioned media from 200 dissociated control or Flag-Pnhd-expressing embryos were incubated with the media from 600 dissociated control embryos, and protein complexes have been pulled down using anti-Flag agarose beads (see Methods) (**Fig. 1A**). After protein separation in an SDS-polyacrylamide gel and Coomassie Blue staining, Flag-Pnhd was visible as prominent 36-38 kD protein bands in the sample from the Pnhd-expressing cells but not in the control sample (**Fig. 1B, C**). Mass spectrometry analyses of the gel slices identified several proteins that preferentially associated with Pnhd (**Fig. 1D**). Among these were Nodal inhibitors, Dand5/Coco/Cerl2 and Lefty/Antivin (Bell et al., 2003; Cheng et al., 2000; Marques et al., 2004; Meno et al., 1998; Montague et al., 2018), and Pnhd itself, consistent with the reported ability of Pnhd to form dimers (Yan et al., 2019). A top candidate (32 identified peptides) corresponded to Admp, a BMP family protein that is expressed in the Spemann organizer and involved in dorsoventral patterning (Moos et al., 1995; Reversade and De Robertis, 2005). These results point to Admp as a candidate endogenous target of Pnhd in *Xenopus* gastrulae.

**Fig. 1.**
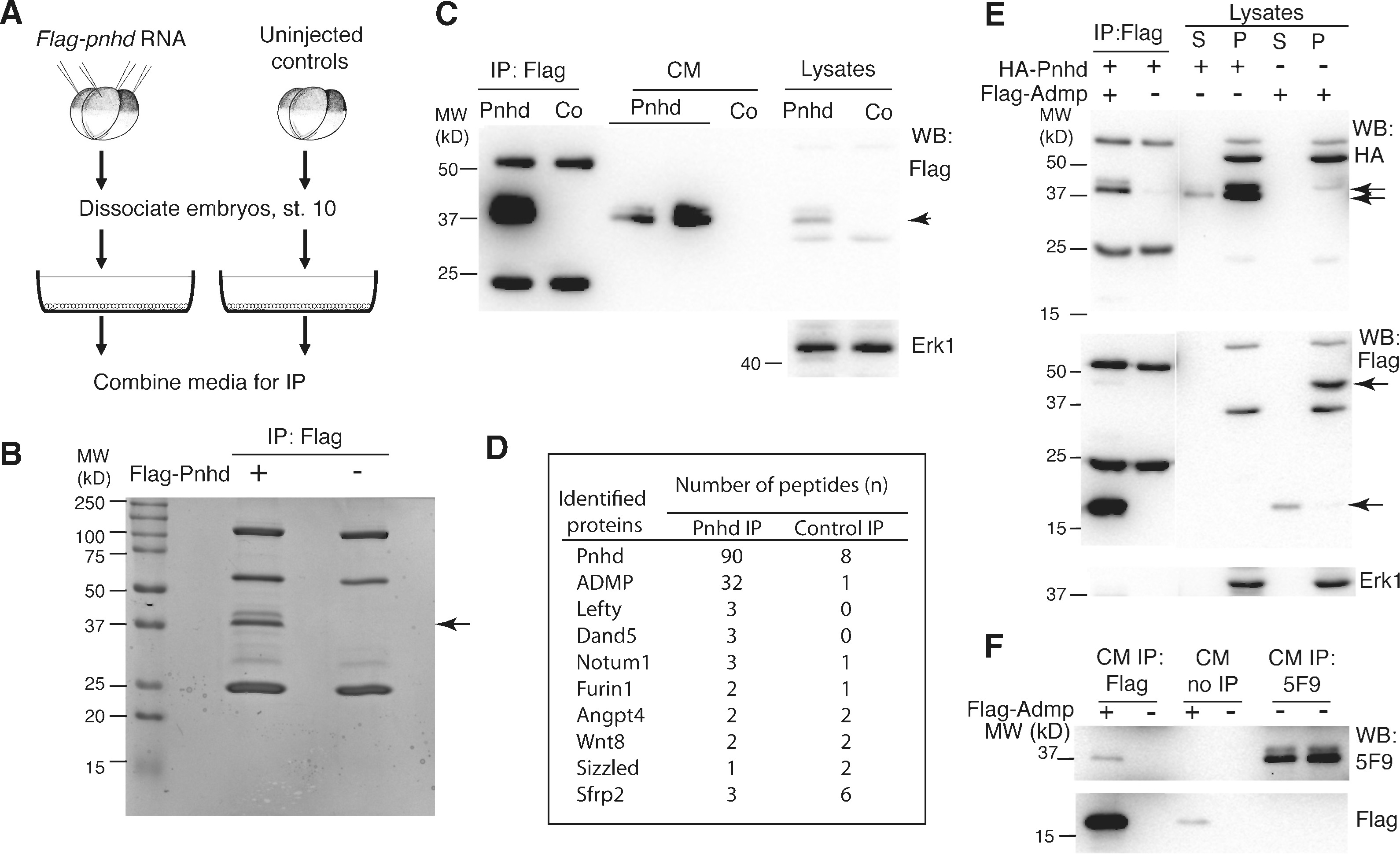
Screening gastrula secretome for Pnhd-associated proteins. A, Experimental scheme. Embryos were injected four times animally with Flag-Pnhd RNA (500 pg) and dissociated at stage 10. After 3 hrs, conditioned media (CM) from dissociated Flag-Pnhd-expressing and control embryos were combined and immunoprecipitated with anti-Flag beads. B. Coomassie blue staining of Flag-Pnhd-containing and control protein pulldowns. Two bands of 36-38 kD correspond to Flag-Pnhd (arrow). C. Immunoblot analysis of immunoprecipitates, CM and cell lysates with anti-Flag antibody. Heavy and light antibody chains are visible in addition to the specific Pnhd bands (arrowhead). Anti-Erk1 antibody serves as loading control for the lysates. D. Numbers of identified peptides for top candidate secreted Pnhd-interacting proteins that were identified by mass spectrometry. E. CM were combined from embryos expressing HA-Pnhd and Flag-Admp as described in (A), and precipitated with anti-Flag antibody. Supernatant (S) or cell pellet (P) fractions from dissociated cell lysates are also shown. Anti-HA antibody recognizes HA-Pnhd as 37-39 kD bands, while anti-Flag antibody detects the unprocessed form of Flag-Admp (45 kD) and mature Flag-Admp (17 kD, arrows). Anti-Erk1 antibody validates the separation of cytoplasmic and secreted proteins. F, Admp binds endogenous Pnhd. CM from dissociated normal embryos (stage 10) or sibling embryos expressing Flag-Pnhd were immunoprecipitated with anti-Flag or anti-Pnhd (5F9) antibodies. After gel separation of protein precipitates, immunoblotting was carried out with the indicated antibodies.

To validate the physical interaction between Pnhd and Admp, we combined the media conditioned by dissociated gastrula cells expressing either HA-Pnhd or Flag-Admp. When Flag-Admp was pulled down by anti-Flag antibodies, HA-Pnhd was readily detected in the precipitate (**Fig. 1E**). No signal was present in the control sample lacking Flag-Admp, indicating that the interaction is specific. This experiment shows that Admp binds to Pnhd.

To confirm this conclusion, we asked whether Admp associates with endogenous Pnhd using Pinhead-specific antibodies (**Suppl. Fig. 1**). The Pnhd protein was easily detected as 36-38 kDa double band in lysates of embryos overexpressing Pnhd but not in normal embryo lysates (**Suppl. Fig. 1A**). This suggests that Pnhd is not abundant but we could visualize endogenous Pnhd by immunoprecipitation from normal gastrula lysates (**Suppl. Fig. 1B**). We found that Pnhd was coprecipitated with Flag-Admp from conditioned media (**Fig. 1F**). Together, these studies indicate that endogenous Pnhd physically associates with Admp.

### Pnhd inhibits Admp signaling but has no effect on BMP4

Admp has a strong ventralizing activity due to its ability to activate BMP receptors and increase Smad1 phosphorylation (Dosch and Niehrs, 2000; Joubin and Stern, 1999; Moos et al., 1995). To assess how Pnhd influences the function of Admp, the corresponding mRNAs were expressed in the marginal zone of four-cell embryos. Consistent with previous reports, *admp* RNA ventralized embryos in a manner similar to that of *bmp4* RNA (**Fig. 2A-C, H)**. *Flag-pnhd* RNA did not cause ventralization on its own at the selected dose **(Fig. 2E, H)**. Strikingly, when *pnhd* and *admp* RNAs were co-expressed, normal dorsal development has been restored (**Fig. 2D, H**). By contrast, BMP4-mediated ventralization was unaffected by Pnhd (**Fig. 2F-H**). These observations demonstrate an inhibitory effect of Pnhd on Admp.

**Fig. 2.**
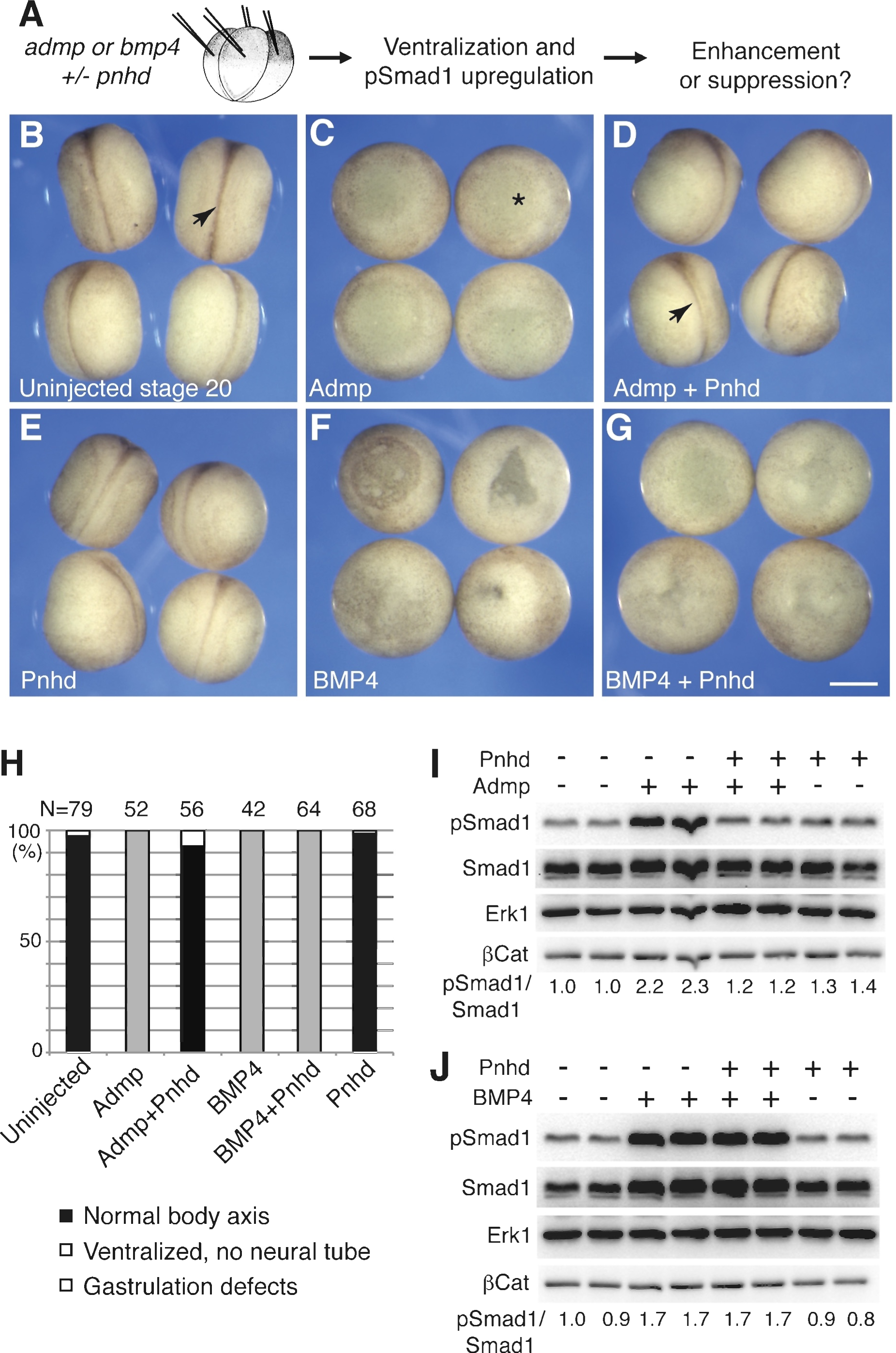
Pnhd inhibits Admp ventralizing activity. A, Experimental scheme. Marginal zone of four-cell embryos was injected with *admp* or *flag-bmp4* RNA (50 pg), and *flag-pnhd* RNA (150 pg) as indicated. B-G, Typical embryo phenotypes at stage 20 are shown. Arrows (B, D) points to neural tubes, asterisk in C marks a ventralized embryo. Bar in G is 400 µm. H, Quantification of the experiments shown in B-G. Ventralization activity in stage 20 embryos has been scored as indicated at the bottom of the panel. Numbers of scored embryos are shown above each bar. Data are representative of three independent experiments. I, J, Pnhd inhibits Smad1 phosphorylation in response to Admp but not BMP4. Embryos were injected as shown in A and harvested at stage 11 for immunoblotting with anti-phospho-Smad1 (pSmad1) and anti-Smad1 antibodies. Normalized ratios of pSmad1 to Smad1 levels are shown. Independent biological replicas are included for each group. Erk1 and β-catenin (βCat) levels control loading.

We next analyzed Smad1 phosphorylation in the injected embryos. Consistent with the negative regulation of Admp, Pnhd inhibited Admp-dependent increase in phospho-Smad1 **(Fig. 2I)**. By contrast, Pnhd had no effect on the stimulation of Smad1 by BMP4 **(Fig. 2J)**. Levels of β-catenin were not altered by Pnhd, Admp and BMP4, suggesting no significant effects of these ligands on Wnt/β-catenin signaling at the doses used (**Fig. 2I, J**). These results confirm that Pnhd is a selective antagonist of Admp. Supporting this conclusion, we found that higher doses of *pnhd* RNA reduced phospho-Smad1 levels in whole embryos **(Suppl. Fig. 2)**.

To further support our observations, we analyzed known Smad1 target genes by RT-qPCR in whole embryos. Whereas Pnhd reversed Admp effects on both dorsal (*chrd1, dkk1)* and ventral (*szl, bambi, ventx*) markers, it did not significantly modulate Bmp4 activity (**Fig. 3A-F)**.

**Fig. 3.**
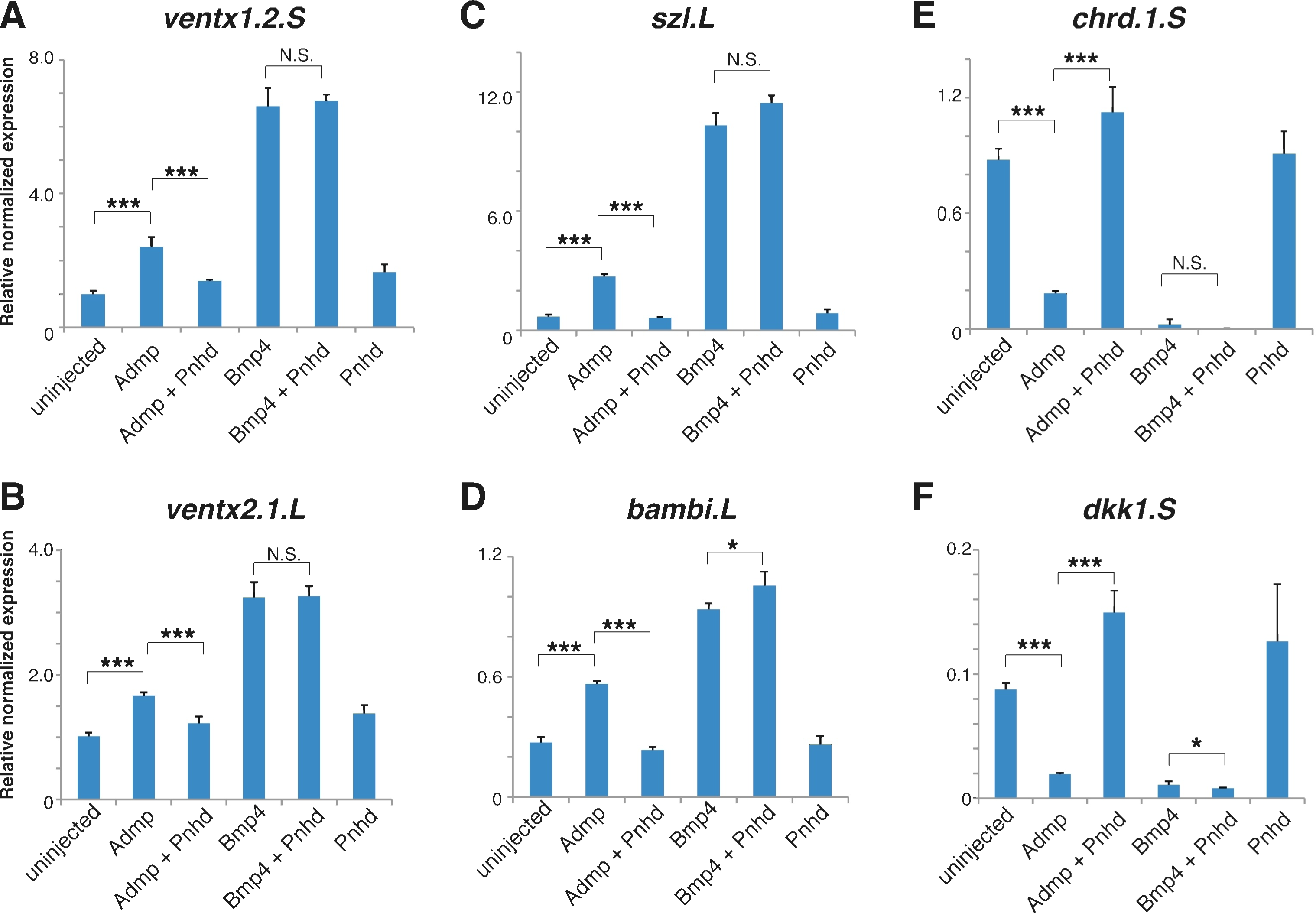
Pnhd regulates Admp-dependent dorsoventral markers. Embryos were injected as described in Fig. 2A legend. RNA was extracted from injected embryos at stage 11.5. RT-qPCR analysis of ventral (*vent1, vent2, szl* and *bambi, A-D*) or dorsal (*chrd1* and *dkk1,* E, F) markers is shown. These data are representative of four experiments. Means +/-s. d. are shown. Significance was determined by the Student’s t-test, p<0.05 (*), p<0.01 (**), p<0.001 (***), p>0.05, non-significant (N.S.).

### Admp activity is enhanced in Pnhd-depleted embryos

*In vivo* Admp activity is likely responsible for only a fraction of total phospho-Smad1 due to the presence of multiple BMP ligands in the early embryo. For this reason, our ability to detect endogenous Pnhd activity may be limited by the insufficient amount of Admp in the embryo. To sensitize the system, we injected embryos with exogenous *admp* RNA and examined Admp-dependent ventralization in *pnhd* morphants. In addition to previously characterized splicing-blocking and translation-blocking morpholino oligonucleotides (MOs) (Ossipova et al., 2020), we designed and validated a non-overlapping splicing-blocking MO, Pnhd-MO2^sp^ (**Suppl. Fig. 3).** Confirming the depletion, we observed a dramatic reduction of Pnhd protein level in embryos injected with Pnhd-MO2^sp^ but not in those injected with a control MO (**Suppl. Fig. 4)**. Using Pnhd-MO2^sp^, we found that the ventralization phenotype caused by *admp* RNA at a suboptimal dose has been significantly enhanced by the coinjection of Pnhd-MO2^sp^ (**Fig. 4A-F)**. A similar effect has been observed for Pnhd-MO^sp^ (**Suppl. Fig. 5A-F)**. These findings demonstrate the inhibitory role of Pnhd in the regulation of Admp activity.

**Fig. 4.**
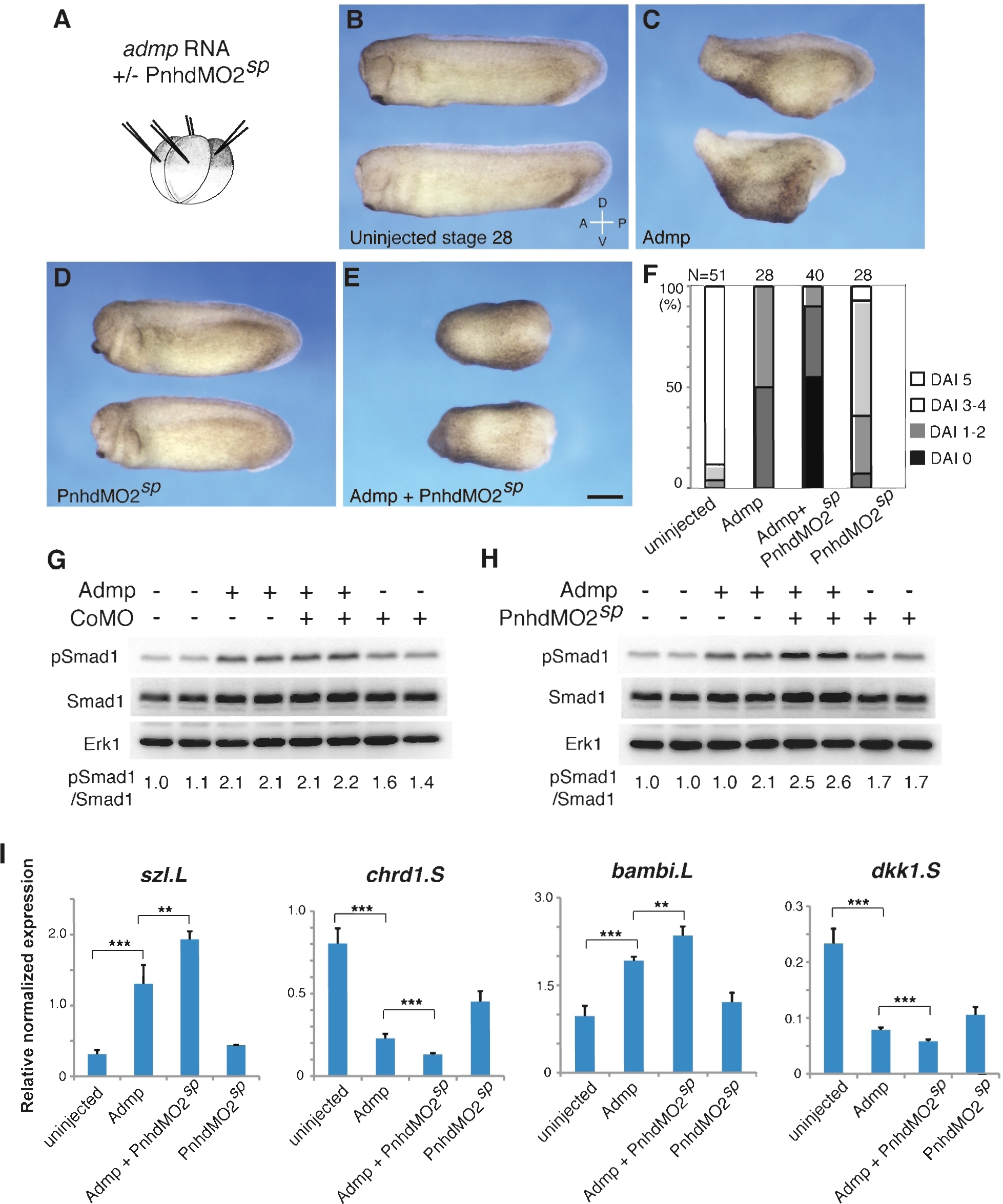
Admp activity is enhanced by Pnhd depletion. A, Experimental scheme. Marginal zone of four-cell embryos was injected with *admp* RNA (50 pg) or PnhdMO2^sp^ (40 ng) as indicated. B-F, Pnhd MO2^sp^ enhanced ventralization caused by Admp. Representative embryos are shown. F. Quantification of the data shown in B-E. Phenotypic scoring was done when uninjected embryos reached stage 28 using dorsoanterior index (DAI). G, H, Immunoblotting of lysates of stage 11 embryos with anti-pSmad1, anti-Smad1 and anti-Erk1 antibodies. G, control MO; H, PnhdMO2^sp^. Normalized ratio of band intensities of pSmad1 relative to total Smad1 is indicated. Erk1 serves as a loading control. Independent biological replicas are included for each group (G, H). Data represent two to three independent experiments. I, The effect of Admp on marginal zone markers is enhanced in Pnhd morphants. RNA was extracted from injected embryos at stage 11.5. RT-qPCR analysis of ventral (*szl* and *bambi*) or dorsal (*chrd1* and *dkk1)* markers is shown. These data are representative of three experiments. Means +/-s. d. are shown. Significance was determined by the Student’s t-test, p<0.01 (**), p<0.001 (***).

Consistent with increased Admp function, we found an upregulation of phospho-Smad1 in Pnhd morphants injected with *admp* RNA. Pnhd-MO2^sp^, Pnhd-MO^sp^ and Pnhd-MO^atg^, but not the control MO, produced this effect (**Fig. 4G, H, Suppl. Fig. 5G, H**). We also observed a small but significant increase in Smad1 phosphorylation in the samples with depleted Pnhd, even without Admp overexpression **(Fig. 4H).** Supporting these observations, the analysis of Smad1-sensitive dorsal (*chrd1, dkk1*) and ventral (*szl, bambi*) gene targets revealed an enhancement of Admp activity in the morphants (**Fig. 4I).**

Taken together, these findings indicate that Pnhd antagonizes Admp to inhibit Smad1 signaling and promote dorsal development.

### Synergistic effects of BMP inhibitors and Pnhd on ectoderm cells

Since both Pnhd and organizer-derived BMP antagonists, such as Chordin, reduce Smad1 signaling in the embryo, we asked whether embryonic cells would exhibit a stronger response to Pnhd in the presence of a BMP inhibitor. Consistent with this hypothesis, the combined inhibition of Admp and BMP ligands caused synergistic phenotypic changes (Reversade and De Robertis, 2005; Willot et al., 2002). To test this prediction, we examined ectodermal explants from the embryos injected with *pnhd* and *chordin* RNAs. By the end of gastrulation, *chordin*-or *pnhd*-expressing ectoderm explants had minimal changes in morphology, however, the explants from the embryos injected with both RNAs dramatically elongated (**Fig. 5A-E, Table 1**). This elongation was clearly detectable as early as the midgastrula stage (**Suppl. Fig. 6**). Importantly, the explant elongation was also stimulated by Pnhd in the presence of a truncated dominant-interfering BMP receptor (Graff et al., 1994; Suzuki et al., 1994) (**Fig. 5F, G**). We conclude that the observed synergy is not due to a unique activity of Chordin, but can be mediated by another Smad1 signaling inhibitor, such as a truncated BMP receptor.

**Fig. 5.**
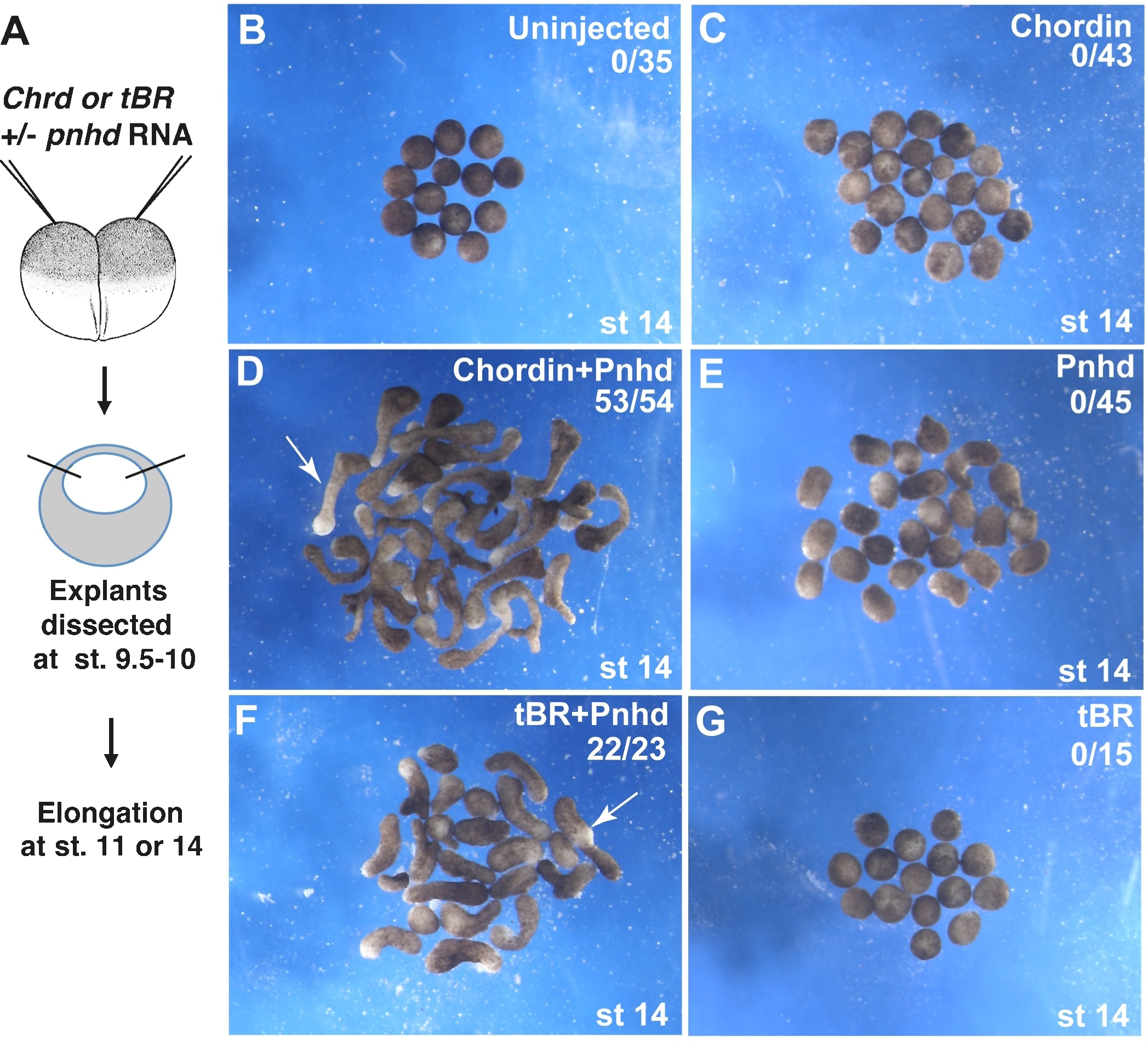
Pnhd cooperates with BMP inhibitors to trigger explant elongation. A, Two-cell stage embryos were injected with *pnhd* (0.5 ng), *chordin* (0.15 ng) or truncated BMP receptor (*tBR*, 0.45 ng) RNA as indicated. Ectoderm explants were dissected at stages 9.5-10 and cultured until stage 14 for morphological examination. B-G, Representative group morphology is shown. Arrows in D and F point to elongating explants. These experiments were repeated 3-7 times.

**Table 1.** Pnhd cooperates with Chordin to promote animal cap elongation.

| RNA injected | Dose | Elongation | Elongation, % | Total N |
| --- | --- | --- | --- | --- |
| Uninjected st 14 | NA | 0/35 | 0 | 35 |
| Chordin | 0.15 ng | 0/43 | 0 | 43 |
| Chordin<br>+Pnhd | 0.15 ng<br>0.5 ng | 53/54 | 98 | 54 |
| Pnhd | 0.5 ng | 0/45 | 0 | 45 |
| tBR<br>+Pnhd | 0.43 ng<br>0.5 ng | 22/23 | 96 | 23 |
| tBR | 0.43 ng | 0/15 | 0 | 15 |
| Uninjected st 11 | NA | 0/21 | 0 | 21 |
| Chordin | 0.15 ng | 0/28 | 0 | 28 |
| Chordin<br>+Pnhd | 0.15 ng<br>0.5 ng | 67/67 | 100 | 67 |
| Pnhd | 0.5 ng | 0/75 | 0 | 75 |
| Uninjected st 14 | NA | 0/39 | 0 | 39 |
| Chordin | 0.15 ng | 0/52 | 0 | 52 |
| Chordin<br>+Pnhd | 0.15 ng<br>0.5 ng | 106/107 | 99 | 107 |
| Pnhd | 0.5 ng | 9/54 | 17 | 54 |
| Uninjected st 15 | NA | 0/13 | 0 | 13 |
| Chordin | 0.15 ng | 0/15 | 0 | 15 |
| Chordin<br>+Pnhd | 0.15 ng<br>0.5 ng | 18/18 | 100 | 18 |
| Pnhd | 0.5 ng | 1/13 | 7.7 | 13 |
| tBR<br>+Pnhd | 0.43 ng<br>0.5 ng | 22/22 | 100 | 22 |
| tBR | 0.43 ng | 0/16 | 0 | 16 |

To examine which gene targets might be responsible for explant elongation, we assessed the synergy of Pnhd and Chordin in target gene activation. RNA sequencing identified 273 genes that were enriched in the animal pole cells expressing both *pnhd* and *chordin* RNAs as compared with the cells expressing *chordin* RNA alone (**Table 2**). The top upregulated genes were notochord-specific markers, including *admp* (Moos et al., 1995) and *shh* (Peyrot et al., 2011), suggesting that the complete inhibition of Smad1 by Chordin and Pnhd promotes dorsal mesoderm. Both *admp* and *shh* are expressed in the organizer at the onset of gastrulation and later in the notochord (Moos et al., 1995; Peyrot et al., 2011). The synergistic activation of *shh, admp,* and *emilin3/emi3,* a *Xenopus* orthologue of a zebrafish notochord-specific gene (Corallo et al., 2013), has been confirmed by RT-qPCR at stage 14 and stage 11, whereas the levels of *sox2* did not change **(Fig. 6A-C**; **Suppl. Fig. 6**). In the absence of BMP inhibitors, Pnhd activated the ventrolateral mesodermal genes, e.g. *cdx4* (**Fig. 6D**), as previously reported (Ossipova et al., 2020). Chordin reduced the induction of these markers, consistent with tissue dorsalization.

**Fig. 6.**
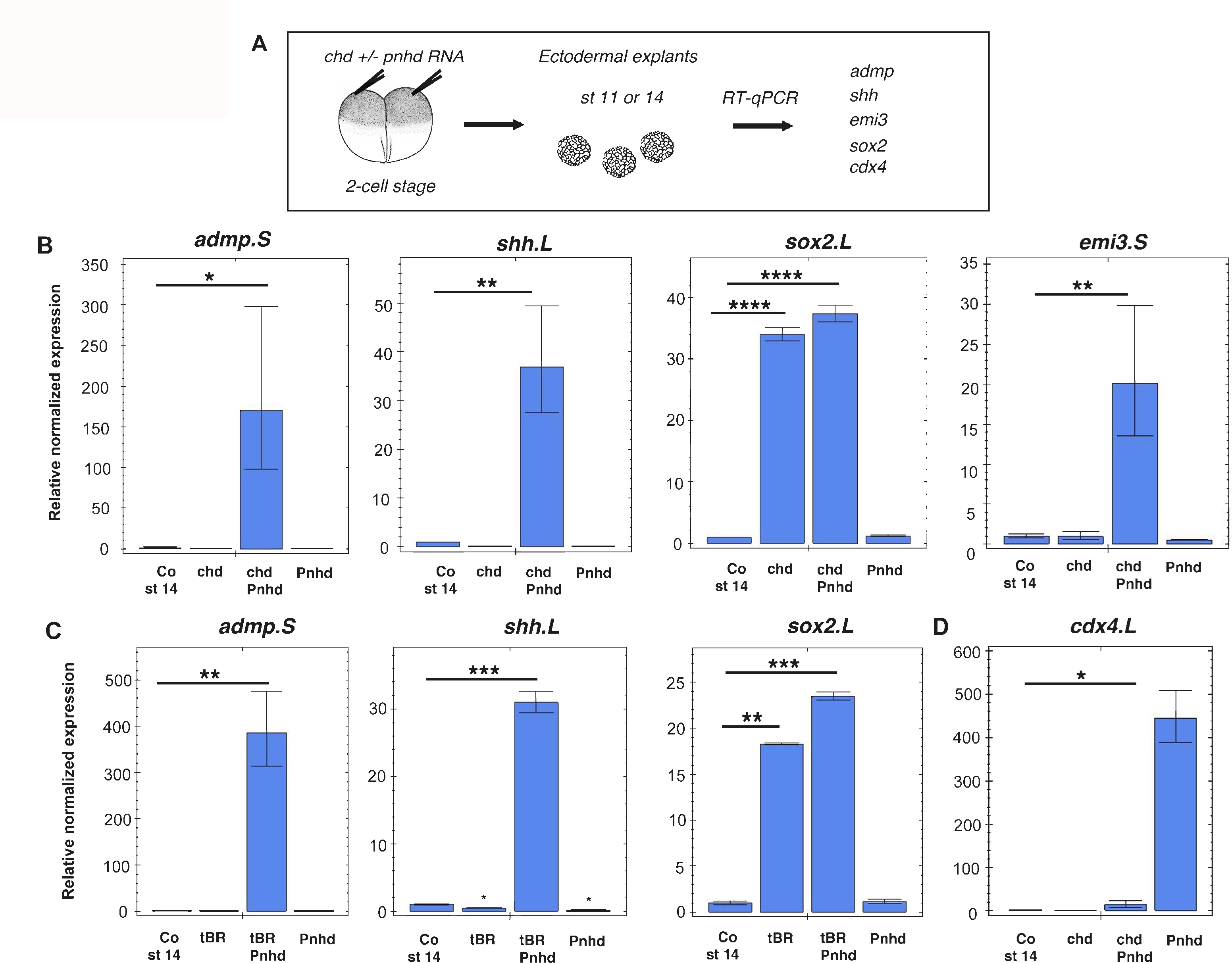
Pnhd synergizes with BMP inhibitors to induce notochord markers. A, Experimental scheme. Two-cell embryos were injected into animal pole region with *pnhd* (0.5 ng) RNA, *chordin* (0.15 ng) or truncated BMP receptor *(tBR*, 0.45 ng) RNA as indicated in B, C. Ectoderm explants were dissected at stages 9.5-10 and cultured until stage 14 to examine notochord markers *admp, shh* and *emi3* by RT-qPCR. B, *Pnhd* synergizes with *Chordin* to induce *admp.S*, *shh.L* and *emi3,* but not *sox2,* at stage 14. C, *Pnhd* synergizes with *tBR* to induce *admp.S* and *shh.L,* but not *sox2*. D, *Chordin* inhibits the induction of *cdx4.L* by *pnhd*. These data are representative of three experiments. Means +/- s. d. are shown. Significance was determined by the Student’s t-test, p<0.05 (*), p<0.01 (**), p<0.001 (***).

**Table 2.**
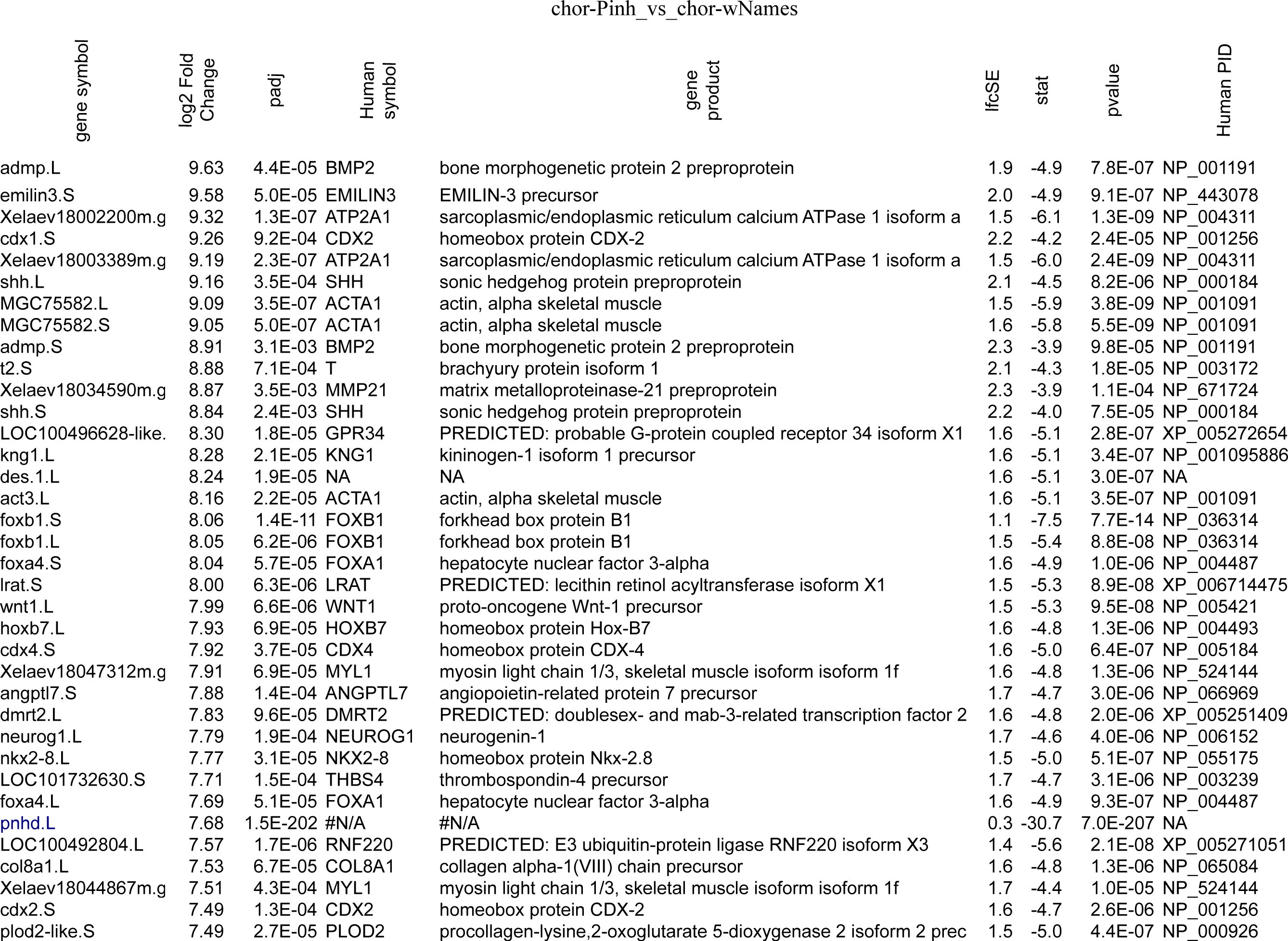
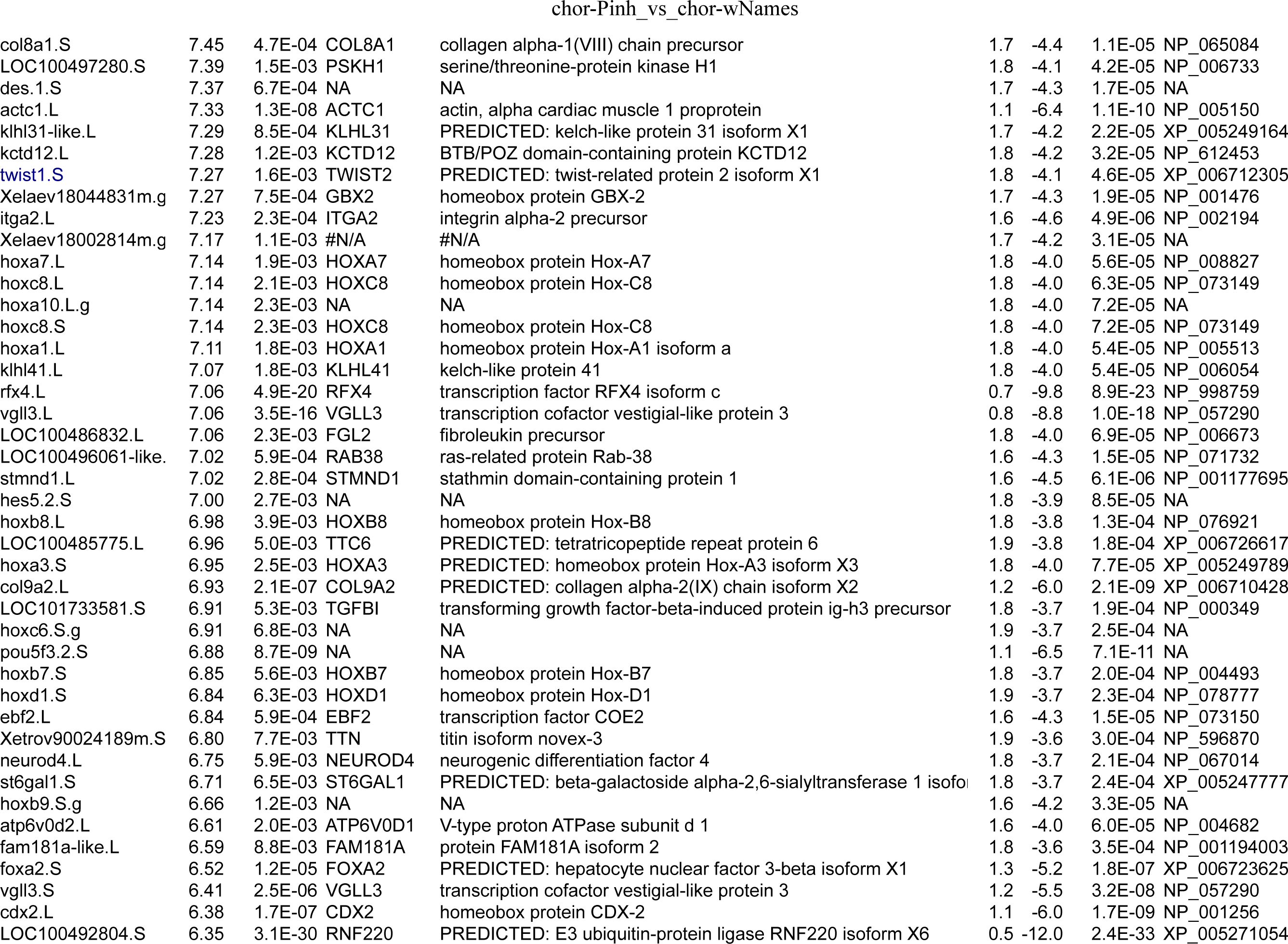
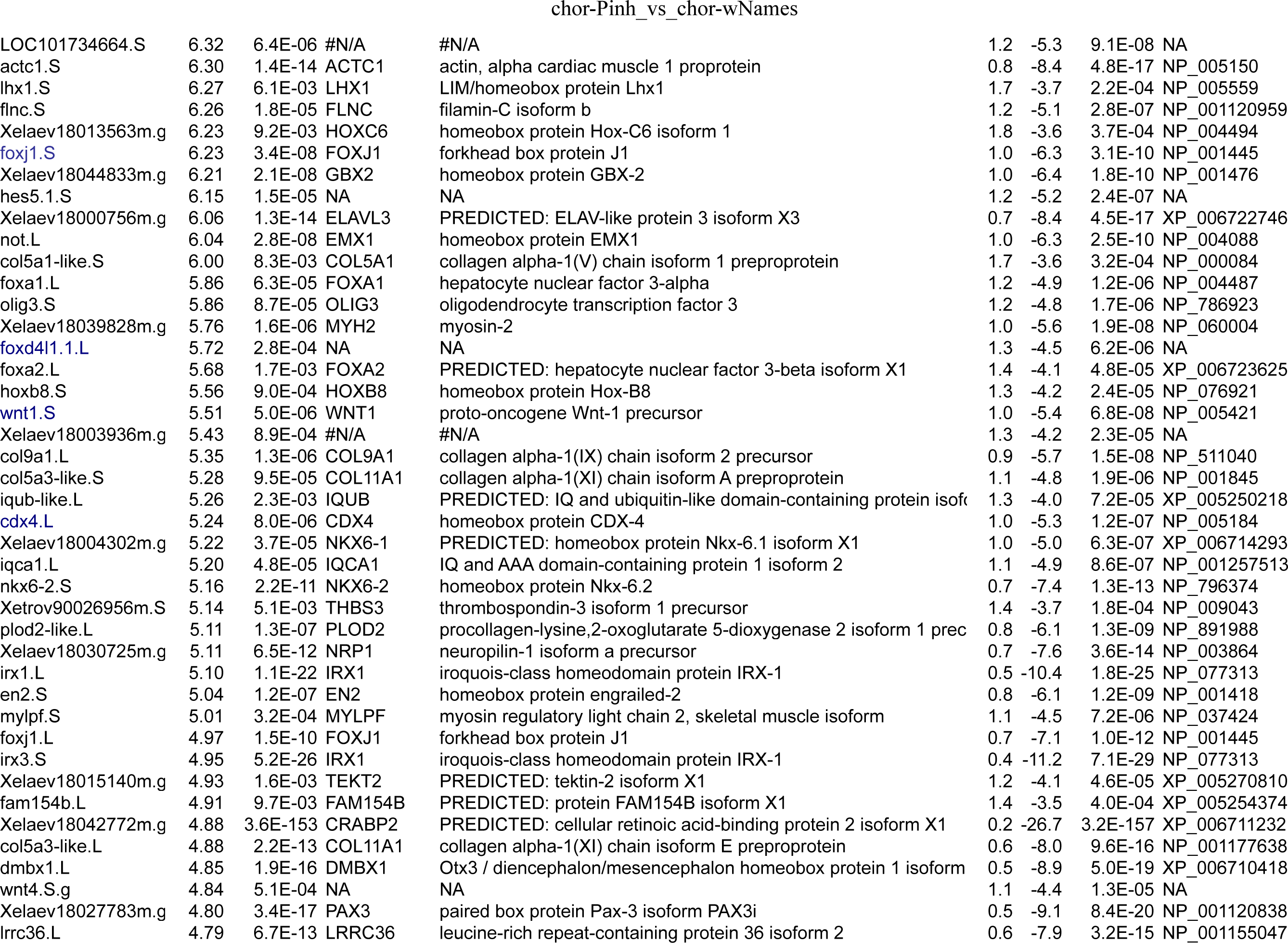
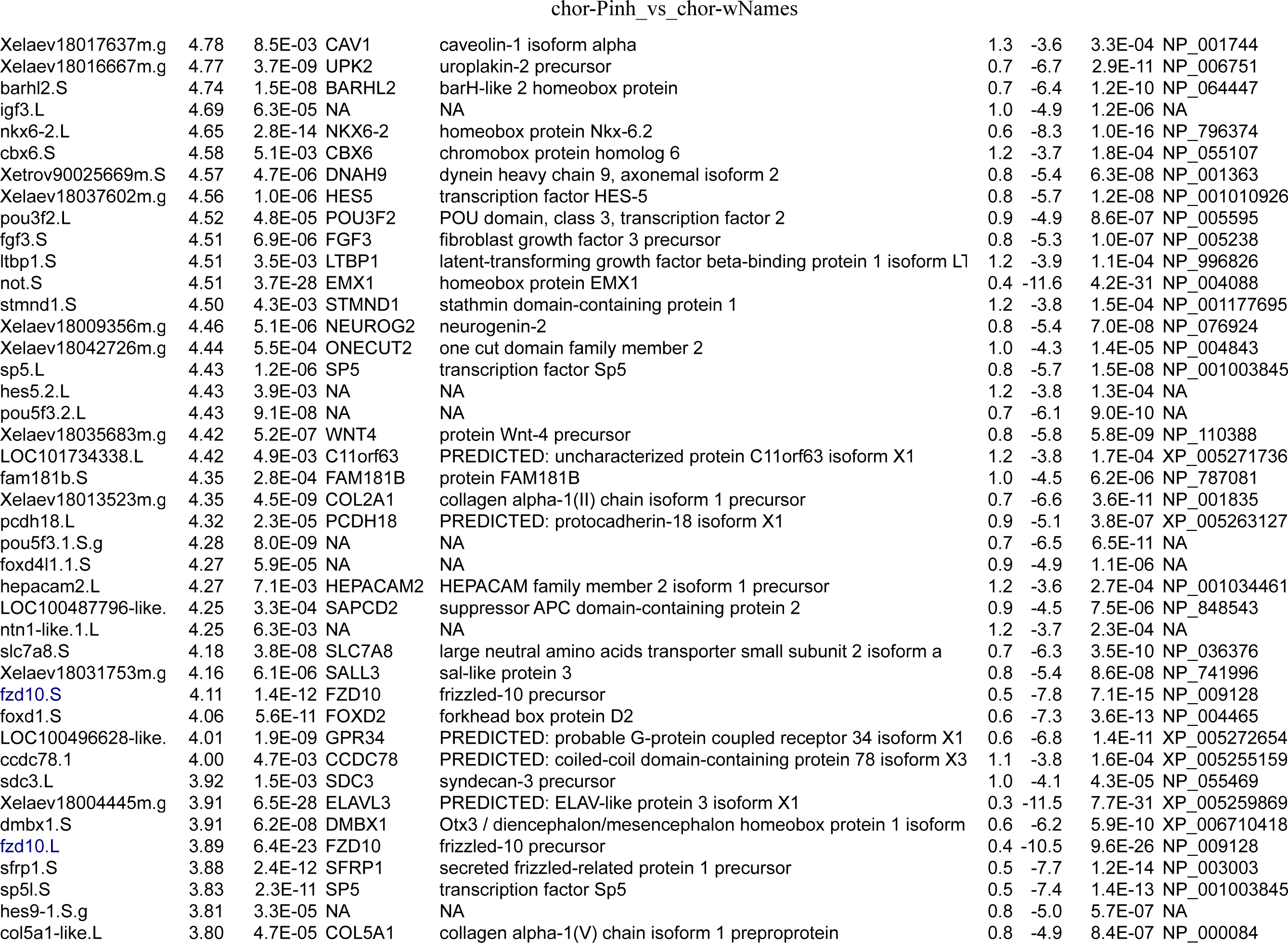
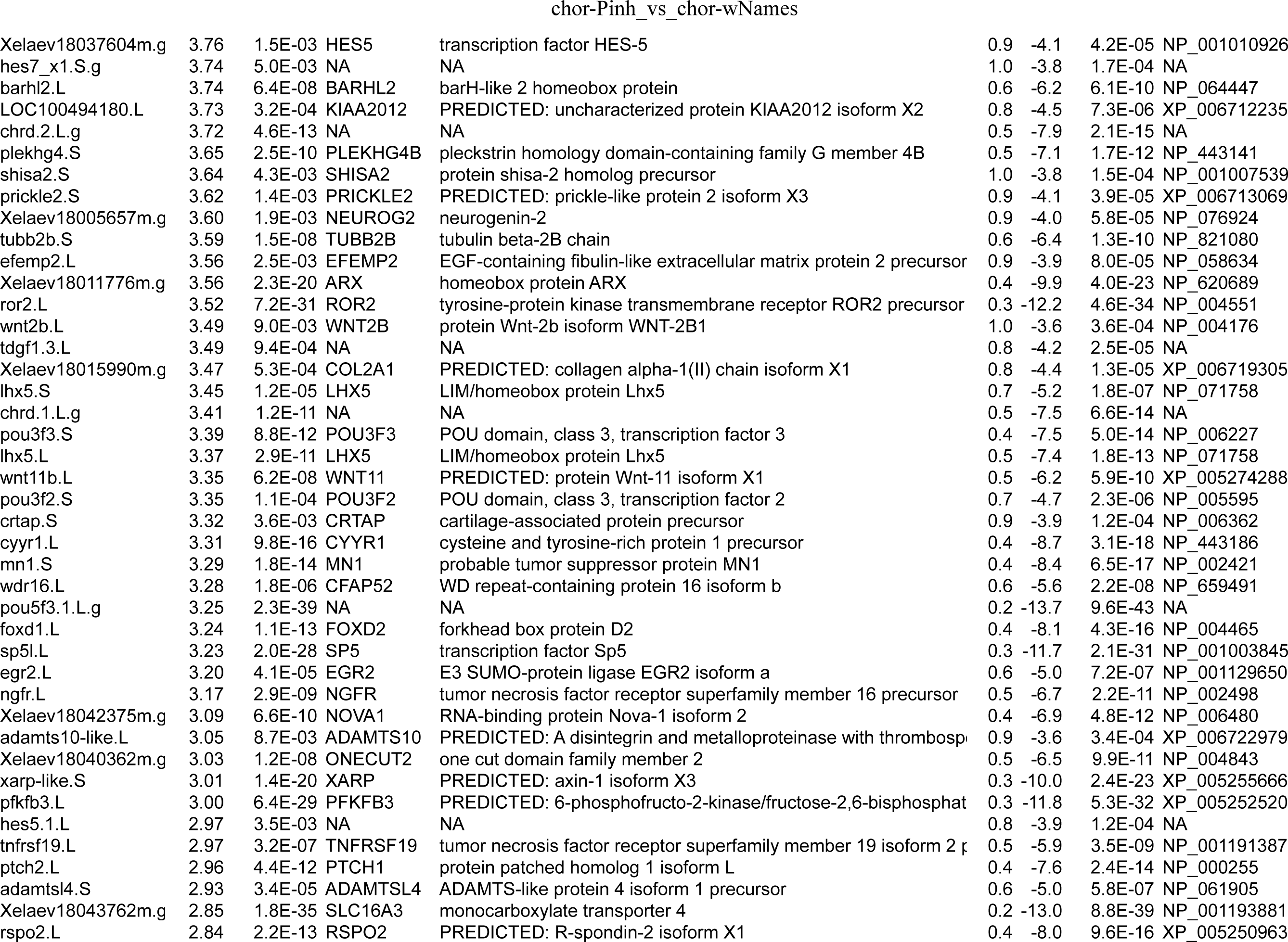
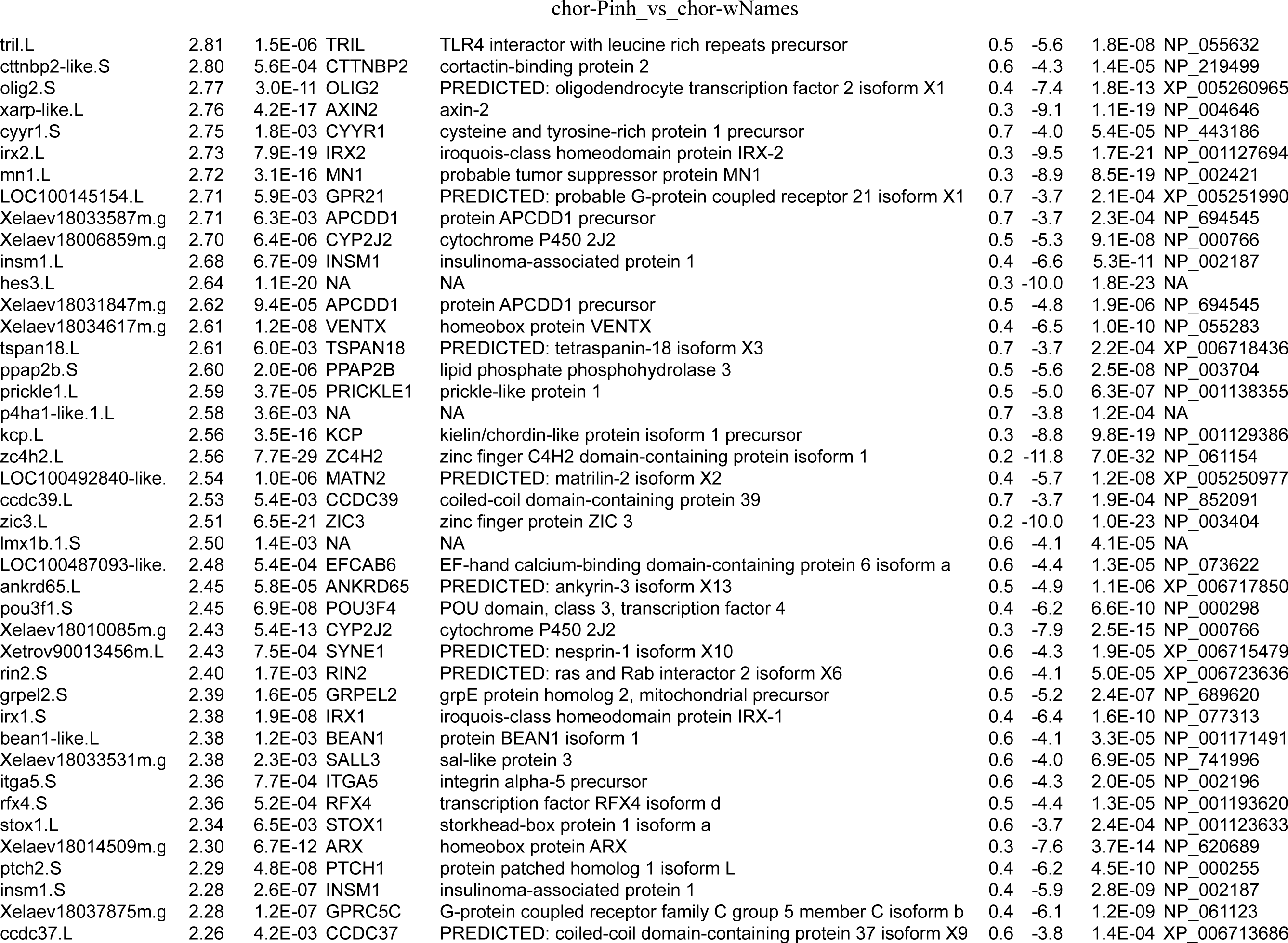
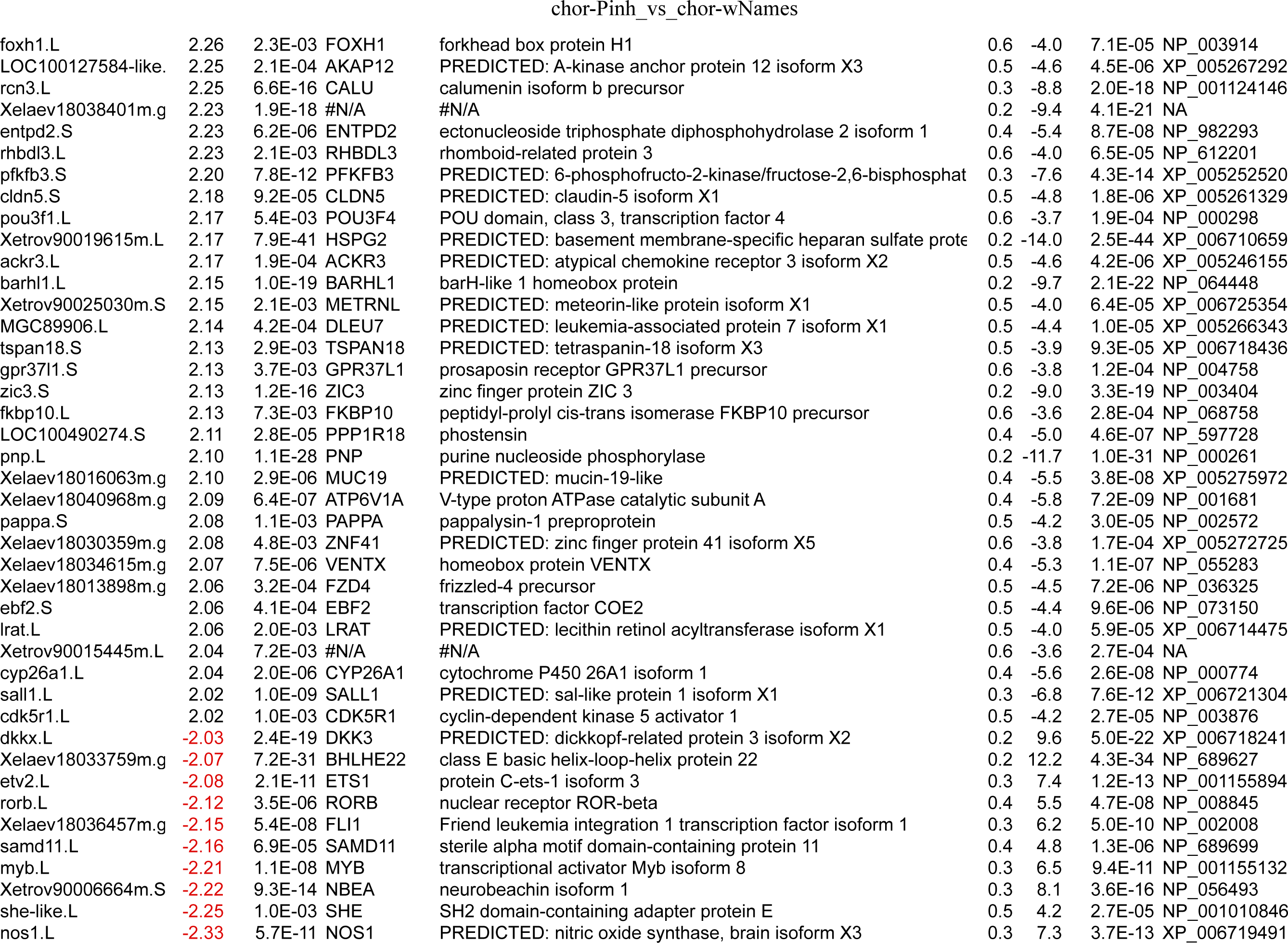
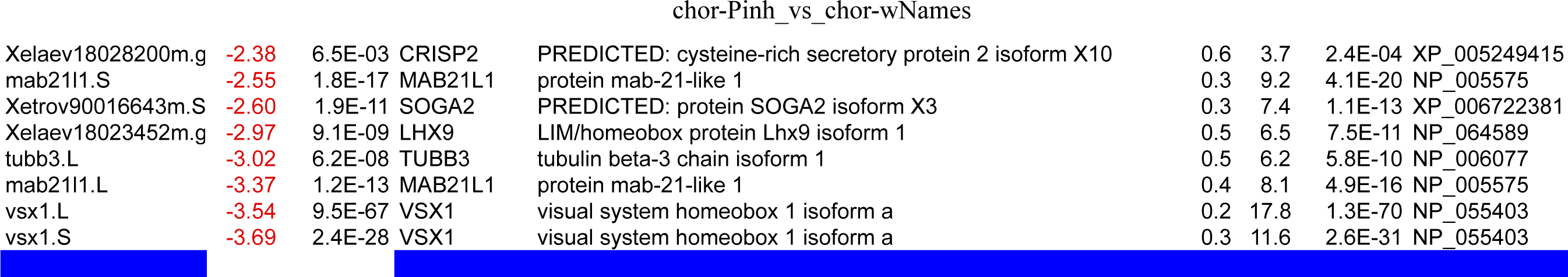
Genes differentially expressed in the ectoderm of embryos injected with *chordin* and *pnhd* RNAs as compared to only *chordin* RNA.

These studies suggest that Pnhd functions together with BMP antagonists to inhibit Smad1 activity and promote dorsal mesoderm formation.

### A requirement for Pnhd in notochord development

To further test the hypothesis that Pnhd promotes dorsal cell fates by antagonizing Admp, we have studied effects of Pnhd depletion in whole embryos by wholemount in situ hybridization with the notochord markers *not* (von Dassow et al., 1993) *and chordin* (Sasai et al., 1994) and the somite marker *myod1* (Hopwood et al., 1989). Embryos were injected with Pnhd-MO^sp^ or Pnhd-MO2^sp^ into four blastomeres of four-cell embryos to target the broad expression of Pnhd in the ventrolateral marginal zone (Ossipova et al., 2020). Embryos injected with either MO revealed consistently narrower and/or weaker expression domains of *chordin* (**Fig. 7A-D**) and *not* (**Fig. 7E-H**), as compared to the uninjected embryos at stage 14. These findings suggest that Pnhd is required for notochord formation.

**Fig. 7.**
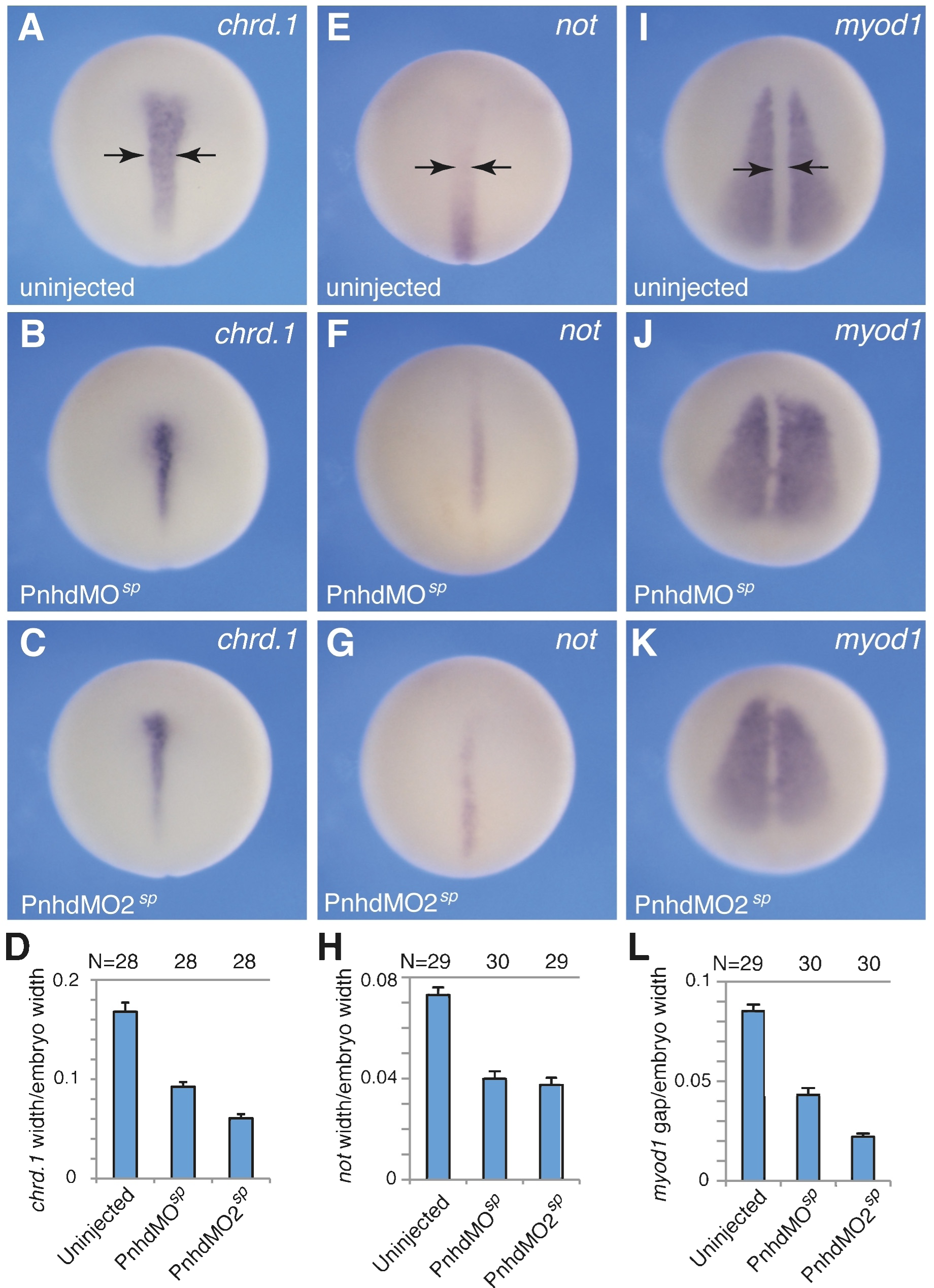
Pnhd is required for notochord specification. A-L, PnhdMO^sp^ or PnhdMO2^sp^ (40 ng each) were injected four times into marginal zone as shown in Fig. 2A. When control embryos reached stage 14, injected or uninjected embryos were fixed for wholemount *in situ* hybridization with the notochord-specific probes *chrd.1* (A-D) and *not* (E-H) or the somitic marker *myod1* (I-L). Dorsal view is shown, anterior is up. Arrows in A, E, I demarcate the width of *chrd.1* or *not* expression domains or the gap in *myod1* expression. D, H, L, Quantification of notochord width (marked by *chrd.1*, *not* or the gap in *myod1)* relative to embryo width. Means +/- s. d. are shown. Number of scored embryos per group are shown above each bar. The data are representative of three independent experiments.

In agreement with these results, the gap between the two *myod1*-expressing domains that corresponds to the notochord was narrower in *pnhd* morphants as compared to the control embryos (**Fig. 7I-L**). Together, these experiments support a role of Pnhd in notochord formation. By contrast, the increased thickness of notochord has been reported for *admp* morphants (Inomata et al., 2013). Thus, we propose that Pnhd regulates notochord size by antagonizing Admp.

## Discussion

Pnhd is expressed in multiple locations in *Xenopus* embryos and interacts with FGF and Nodal signaling during early stages of ventroposterior mesoderm specification (Ossipova et al., 2020). In this study, we have used mass spectrometry to analyze Pnhd-associated proteins pulled down from the medium conditioned by dissociated *Xenopus* gastrula cells (the gastrula secretome). We identified several candidate secreted proteins that appear to physically associate with Pnhd. The top hit in this screen was Admp, a member of the BMP family. Our biochemical and functional studies confirmed the specific binding of Admp by Pnhd and revealed that Pnhd inhibits Admp activity both phenotypically and molecularly by modulating phospho-Smad1 level. By inactivating Admp, Pnhd contributes to the specification of dorsal mesoderm including the notochord at later stages (**Fig. 8**). Thus, Pnhd might play multiple roles by interacting with distinct signaling pathways at different developmental stages.

**Fig. 8.**
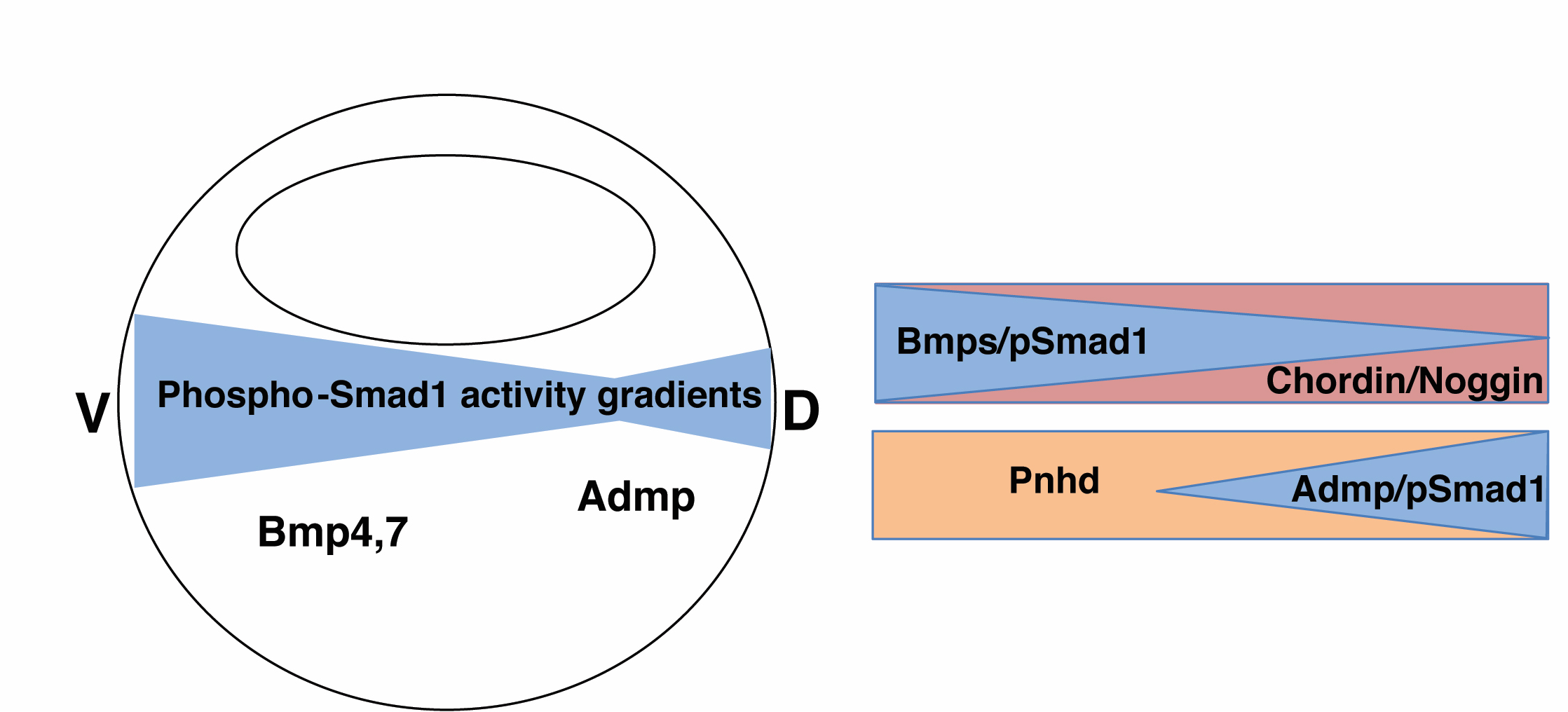
A model of Pnhd effects on dorsal mesoderm, Admp and Smad1 activity. Major activators of Smad1 phosphorylation in the *Xenopus* gastrula are ventrolaterally-expressed Bmp4/7 and dorsal Admp. *Pnhd* transcripts are distributed in the ventrolateral marginal zone, complementary to the dorsal expression of *admp*. Dorsal organizer signals, such as Chordin, reduce Smad1 phosphorylation triggered by Bmp4/7. A similar antagonistic effect of Pnhd on Admp is proposed to further modulate Smad1 activity gradients.

Notably, *pnhd* and *admp* genes are positioned next to each other in the genome of lower vertebrates and birds, but have been eliminated from the mammalian genomes suggesting coevolution (Imai et al., 2012). While the physical binding between Pnhd and Admp has been already reported in *Ciona*, it remained unclear whether and how Pnhd influences Admp (Imai et al., 2012). Zebrafish *pnhd* mutant embryos do not have a strong morphological phenotype on their own (Yan et al., 2019). Nevertheless, in the zebrafish model, Pnhd was shown to activate BMP receptors and act redundantly with Admp (Yan et al., 2019), These observations indicate that the interaction between Pnhd and Admp may be context-dependent. Our results suggest that Pnhd could directly sequester Admp, potentially interfering with the binding of BMP receptors and Smad1 phosphorylation. In support of this model, *Xenopus* Pnhd morphants are microcephalic (Kenwrick et al., 2004; Ossipova et al., 2020) and have a narrow notochord (this study), whereas Admp inhibition causes enlarged head structures and widened notochord (Inomata et al., 2013; Inui et al., 2012; Reversade and De Robertis, 2005). Alternatively, Pnhd might form heterodimers with Admp to prevent its signaling. This possibility is less likely, because Pnhd is structurally very different from BMPs, with multiple cystine knot domains. Unlike BMPs, Pnhd does not appear to undergo Furin-dependent proteolytic processing, because the proteins tagged at the N- and C-termini have the same mobility.

The *pnhd* and *admp* genes have been evolutionarily conserved and expressed in mutually exclusive domains in *Ciona*, zebrafish and *Xenopus* embryos, suggesting feedback regulation (Imai et al., 2012). Although Pnhd is produced in the ventrolateral marginal zone (Kenwrick et al., 2004; Ossipova et al., 2020), and Admp is in the dorsal organizer, we find that *admp* transcription is strongly upregulated by Pnhd in the presence of Chordin. Whereas this observation is at odds with the transcriptional repression predicted by enhancer analysis in *Ciona* (Imai et al., 2012), we hypothesize that Chordin dorsalizes the ventrolateral mesoderm induced by Pnhd to activate several notochordal markers, including *admp*. The induction of *admp* might be particularly strong because of the compensatory feedback needed to restore Admp levels after its inactivation by Pnhd. Further analysis of the admp gene regulatory network is necessary to fully understand our observations.

The finding that Pnhd antagonizes Admp reveals an additional level of regulation of Smad1 signaling activity. This activity is primarily controlled by ventral BMP ligands and their dorsal antagonists Chordin and Noggin (De Robertis and Kuroda, 2004; Tuazon and Mullins, 2015). Other secreted molecules, such, as Admp, Tolloid-like proteases and Sizzled/Ogon (Lee et al., 2006; Muraoka et al., 2006) also contribute to Smad1 regulation. Dorsal stimulation of Smad1 signaling by Admp has been proposed to control embryo scaling (Ben-Zvi et al., 2008; Inomata et al., 2013; Reversade and De Robertis, 2005). Our study further extends these observations to indicate that the interaction of Pnhd and Admp is critical for the refining of Smad1 signaling activity during dorsoventral patterning.

## Limitations of the Study

We have shown that *pnhd* is necessary to fine tune Smad1 signaling by antagonizing Admp. Although the mutants of Admp and Pnhd can be generated, it is not practical to pursue classical genetic analysis using *Xenopus laevis*, due to the long generation time. Moreover, the exact molecular mechanism underlying the observed interaction remains unclear. Additional studies are needed to determine whether Pnhd sequesters Admp to interfere with Admp binding to BMP receptors or promotes the interaction of Admp with other secreted inhibitors such as Chordin.

## STAR Methods

### Resource Availability

*Lead Contact* Futher information and requests for reagents should be directed to and will be fulfilled by the Lead Contact, Sergei Sokol.

### Materials availability

All data generated in this study are included in this published article and its supplemental information.

### Data and Code Availability

The RNA sequencing datasets generated during this study are available at the NIH GEO (submission number GSE168370). All other data produced in this study are included into this published article.

### Experimental Model and Subject Details

Mature female and male Xenopus laevis frogs were obtained from NASCO (Fort Atkinson, WI). Animals were maintained in a recirculating tank system with regularly monitored temperature and water quality (pH, nitrate and nitrite levels) Xenopus laevis were housed at a temperature of 18-20C. Frogs were fed with food pellets (NASCO, cat#SA05960). All experimental protocols involving frogs were performed in strict accordance with the recommendations in the Guide for the Care and Use of Laboratory Animals of the National Institutes of Health. The protocol 04-1295 was approved by the IACUC of the Icahn School of Medicine at Mount Sinai.

### Method Details

All methods can be found in the accompanying Transparent Methods supplemental file.

#### Plasmids, in vitro RNA synthesis and morpholino oligonucleotides

pCS2-Flag-Pnhd and pCS2-HA-Pnhd plasmids have been generated by PCR from the *X. laevis* DNA clone for *pnhd.L* (accession number NM_001127751) obtained from Dharmacon. Flag or HA tags were introduced after the predicted signal peptide sequence of Pnhd. pSP64T3-Flag-Admp is generated by PCR from *Xenopus laevis admp.S* cDNA obtained from Bill Smith (Kumano et al., 2006). pCS2-Chrodin was from E. De Robertis (Sasai et al., 1995). Flag tag was introduced in Admp by PCR-based mutagenesis, four amino acids downstream of the conserved processing site RLGR.

Capped mRNAs were synthesized using Ambion mMessage mMachine kit (ThermoFisher). pCS2-Flag-Pnhd, pCS2-HA-Pnhd, pCS2-Pnhd-Flag, pCS2-Pnhd, pSP64T3-Flag-Admp, pSP64T3-Admp, pCS2+Chordin (obtained from Eddy De Robertis), pCS2-Flag-BMP4 (obtained from Andrei Zaraisky, (Bayramov et al., 2011)) and pSP64T-tBR (Graff et al., 1994) were used. Splicing blocking MOs, Pnhd MO^sp^, 5’-CCTGTTCATCACGCTACCATCTAAA-3’ and Pnhd-MO2^sp^, 5’-GGACTACCAGAGATATCTGTAATAA-3’, translation blocking Pnhd MO^atg^, 5’-ACAAGAAAAGATGTTCCATGTCTG-3’; and control MO (CoMO), 5’-GCTTCAGCTAGTGACACATGCAT-3’, were purchased from Gene Tools (Philomath, OR).

#### Xenopus embryo culture, microinjections, and production of secreted proteins

*In vitro* fertilization and culture of *Xenopus laevis* embryos were carried out as previously described (Dollar et al., 2005). Staging was according to Nieuwkoop and Faber (Nieuwkoop and Faber, 1967). For microinjections, 2 to 4-cell embryos were transferred into 3 % Ficoll in 0.5x Marc’s Modified Ringer’s (MMR) solution (50 mM NaCl, 1 mM KCl, 1 mM CaCl_2_, 0.5 mM MgCl_2_, 2.5 mM HEPES pH 7.4) (Peng, 1991) and 10 nl of mRNA or MO solution were injected into two blastomeres of 2 cell embryos or four blastomeres of 4 cell embryos either animally or sub-equatorially. Injected embryos were transferred into 0.1 MMR at blastula stages. Amounts of injected mRNA per embryo have been optimized in preliminary dose-response experiments and are indicated in figure legends.

Secreted proteins were produced after dissociating whole embryos in Ca/Mg-free medium essentially as described (Eroshkin et al., 2016). At early gastrula stage, vitelline membrane was removed and 60 embryos were transferred into 600 µl of Ca/Mg-free medium (88 mM NaCl, 1 mM KCl, 2.4 mM NaHCO_3_, 7.5 mM Tris-HCl, pH 7.6) per well of 4 well dish. After shaking at 80-100 rpm for 3 hrs (with 2 mM EDTA for the last hour), the medium (500 µl) was collected from each well and cleared by centrifugation at 100 g for 1 min. For immunoprecipitation, 4.5 mM CaCl_2_, 1.6 mM MgCl_2_ and 1 mM phenylmethylsulfonylfluoride (PMSF, Sigma) were added. The remaining cell pellets were lysed in 500 µl of the lysis buffer (50 mM Tris-HCl pH 7.6, 50 mM NaCl, 1 mM EDTA, 1% Triton X-100, 10 mM NaF, 1 mM Na_3_VO_4_, 1 mM PMSF) and the lysates were collected after centrifugation for 4 min at 16000 g for immunoblotting.

For whole embryo lysates, 5 embryos were lysed in 85 μl of the lysis buffer and the lysates were collected after the same centrifugation. For animal cap experiments, both blastomeres of the 2-cell embryo were injected in the animal pole region. Ectoderm explants were prepared at stages 9+ to 10, and cultured in 0.6xMMR in the presence of gentamicin (10 µg/mL) until the indicated time. Embryo phenotypes at stage 28 were quantified according to the dorso-anterior index (DAI)(Kao and Elinson, 1988). DAI 5, normal embryos; DAI 4, small head with eyes and cement glands; DAI 3, small head with cement glands; DAI 2, rudimentary head without eyes nor cement glands; DAI 1, posterior body axis lacking head; DAI 0, completely ventralized.

#### Immunoprecipitation, western blot analysis and blue staining

For immunoprecipitation of Flag-Pnhd protein in mass spectrometry experiments, 1.5 ml of conditioned medium from 200 Flag-Pnhd-expressing or control embryos were combined with 5 ml of conditioned medium from 600 control embryos and incubated with 25 µl of anti-FLAG agarose beads (Sigma) at 4°C overnight. For coprecipitation of Flag-Admp and HA-Pnhd, the HA-Pnhd- and Flag-Admp-containing or control conditioned media (500 µl each) were combined and incubated with 2 µl of anti-Flag agarose beads at 4° C overnight. For endogenous Pnhd, the conditioned medium (500 μl) from embryos injected with PnhdMO2^sp^ or control MO (40 ng x 4) was precipitated with 20 μl of the supernatant containing anti-Pnhd 5F9 antibodies (a gift of D. Alfandari), followed by incubation of 4 μl of Protein A sepharose beads. The beads were washed with phosphate buffered saline (PBS) and boiled in the sample buffer. For immunoblotting, 3 or 10 μl of immunoprecipitated proteins, 10 μl of conditioned medium or dissociated cell lysates or 10 μl of embryo lysates were separated by SDS-PAGE. Immunoblotting was carried out essentially as described (Itoh et al., 2005). For Coomassie blue staining, 15 μl of immunoprecipitated material (25 μl total) was loaded and separated on a gel followed by SDS-PAGE and the gel was stained with Simply Blue (Pierce) according to the manufacturer’s protocol. Gel slices were excised after Simple Blue staining and subjected to medium gradient LC-MS/MS analysis carried out by the Keck Proteomics Laboratory at Yale University.

The following primary antibodies were used: mouse anti-FLAG (M2, Sigma), mouse anti-HA (12CA5), anti-Myc (9E10), rabbit anti-Erk1 (K23, Santa Cruz), anti-pSmad1/5 S463/465 (41D10, Cell Signaling Technology), anti-Smad1 (Invitrogen), rabbit anti-β-catenin (Invitrogen) and anti-Pnhd antibodies (5F9, 8G12, 11D5; gifts of D. Alfandari). The detection was carried out by enhanced chemiluminescence as described (Itoh et al., 2005), using the ChemiDoc MP imager (BioRad). Normalized ratios of band intensities of pSmad1 to Smad1 were calculated using NIH imageJ software.

#### Ectoderm explants, RNA sequencing, RT-PCR and qRT-PCR

Embryos were injected with various MOs and RNAs as indicated in figure legends. Ectoderm explants were prepared and cultured until the required stage. RNA was extracted from 15-30 animal pole explants or 3-4 whole embryos using the RNeasy kit (Qiagen). For RT-PCR, cDNA was made from 1-2 µg of total RNA using the first strand cDNA synthesis kit (Invitrogen) or iScript (Bio-Rad) according to the manufacturer’s instructions.

For RNA sequencing, ectoderm explants expressing *chordin* RNA (150 pg), *pnhd* RNA (0.5 ng) or both, and control uninjected explants were cultured until stage 13/14. cDNA library preparation and paired-end 150 bp sequencing were performed by Novogene (Sacramento, CA) using Illumina HiSeq2000 analyzers. The raw reads (FASTQ files) were filtered to remove reads containing adapters or reads of low quality. The sequences were mapped to the *Xenopus* genome version XL-9.1_v1.8.3.2 available at http://www.xenbase.org/other/static/ftpDatafiles.jsp using the software hisat2 (Kim et al., 2015). The differentially expressed genes (DEGs) were detected using DESeq (Anders and Huber, 2010) with the two-fold change cutoff. The p-value estimation was based on the negative binomial distribution, using the Benjamini-Hochberg estimation model with the adjusted p < 0.05. *Pnhd* and *chordin*-induced DEGs have been assessed using two independent samples in the same RNA sequencing experiment. The entire dataset has been submitted to the NIH GEO (submission number GSE168370).

For RT-qPCR, the reactions were amplified using a CFX96 light cycler (Bio-Rad) with Universal SYBR Green Supermix (Bio-Rad). Primer sequences used for RT-PCR and RT-qPCR are shown in **Table 3**. The reaction mixture consisted of 1X Power SYBR Green PCR Master Mix, 0.3 µM primers, and 1 µl of cDNA in a total volume of 10 µl.

**Table 3.** Primers used for RT-qPCR and RT-PCR.

| **Gene name** | **Forward primer** | **Reverse primer** |
| --- | --- | --- |
| ef1a.1.S | 5'-acc ctc ctc ttg gtc gtt tt -3', | 5'-ttt ggt ttt cgc tgc ttt ct -3' |
| admp.S | 5'-atg tat act gtc tga aag gg -3' | 5'-ggt ggc tat aaa tga tac tt -3' |
| emilin3 | 5'- ggg cac cag tgg gga act gc -3' | 5'- tc cca cgc agc agg tct gtc -3' |
| cdx4.L | 5'- tga ttt atc acc taa cca g -3' | 5'- gtc cca gat gga tga gga ga -3' |
| shh.L | 5'- aac aca cct ggg cac acc tc -3' | 5'- tcc aaa agc caa gtc cct at -3' |
| sox2.L | 5'-tca cct ctt ctt ccc att cg -3' | 5- cga cat gtg cag tct gct tt -3' |
| ventx1.2.S | 5'-ttc cct tca gca tgg ttc aac-3' | 5'- gca tct cct tgg cat att tgg-3' |
| ventx2.1.L | 5'-tga gac ttg ggc act gtc tg-3' | 5'-cct ctg ttg aat ggc ttg ct-3' |
| szl.L/S | 5'- gtc ttc ctg ctc ctc tgc-3' | 5'- aac agg gag cac agg aag-3' |
| chrd1.S | 5'-cat gct ctt tcg aag gtc aa-3' | 5'- gat cac aaa tca cgg tac gc-3' |
| bambi.L | 5'- gca caa tgg tcc agt agg gg-3' | 5'-ttc gtc ctc cta aag atc ag-3' |
| dkk1.S | 5'-aca gag cct gaa gga cct tc-3' | 5'-gtt cag gga aga cca gag ca-3' |

***Primers used for RT-PCR***
| **Gene name** | **Forward primer** | **Reverse primer** |
| --- | --- | --- |
| ef1a | 5'- cag att ggt gct gga tat gc -3' | 5- act gcc ttg atg act cct ag -3' |
| pnhd.L | 5'- tga tga aca ggc aga atg ac-3' | 5- gat tca aag gga gaa tct gg -3' |

The ΔΔCT method was used to quantify the results. All samples were normalized to control uninjected embryos or explants. Transcripts for *elongation factor 1a1 (ef1a1*) were used for normalization. Data are representative of two to three independent experiments and shown as means +/- standard deviation. Statistical significance was assessed by two-tailed Student’s *t*-test (\**p*<0.05, \*\**p*<0.01, \*\*\**p*<0.001 and \*\*\*\**p*<0.0001).

#### Wholemount in situ hybridization

*Wholemount in situ* hybridization (WISH) was carried out as described (Harland, 1991) and pigments were bleached as described (Mayor et al., 1995). Diogoxigenin-rUTP– labeled RNA probes were prepared by in vitro transcription of linearized plasmids of pBS-*chrd.1* (Sasai et al., 1994), pBSKS*-not* (von Dassow et al., 1993), and pBSSK*-myod1* (Hopwood et al., 1989) with T3 or T7 RNA polymerases and the RNA labeling mix containing digoxigenin-rUTP (Roche). All data are representative of two to three independent experiments. Measurement of width of expression domains of *chrd.1* and *not* or expression gap of *myod1* as well as embryo width was performed with NIH ImageJ software.

### Quantification and statistical analysis

Each figure legend contains quantification details, including frequencies of embryonic phenotypes or numbers of explants examined (n), the mean values, and the s.d. In addition, information about the statistical tests used for measuring significance and interpretation of p values is provided. Statistical analyses and graphic results for RT-qPCR were generated using the CFX Maestro software (Biorad).

## Acknowledgement

We thank Eddy DeRobertis, Bill Smith, David Kimelman, Jonathan Graff, Andrei Zaraisky and John Gurdon for plasmids, Dominique Alfandari for the anti-Pnhd antibodies, Aurelian Radu for his help with RNA sequencing data analysis, and members of the Sokol laboratory for discussions. This study was supported by the following NIH grants: R01HD092990, R01GM116657, R24OD21485.

## Supplementary Figure legends

**Suppl. Fig. 1.**
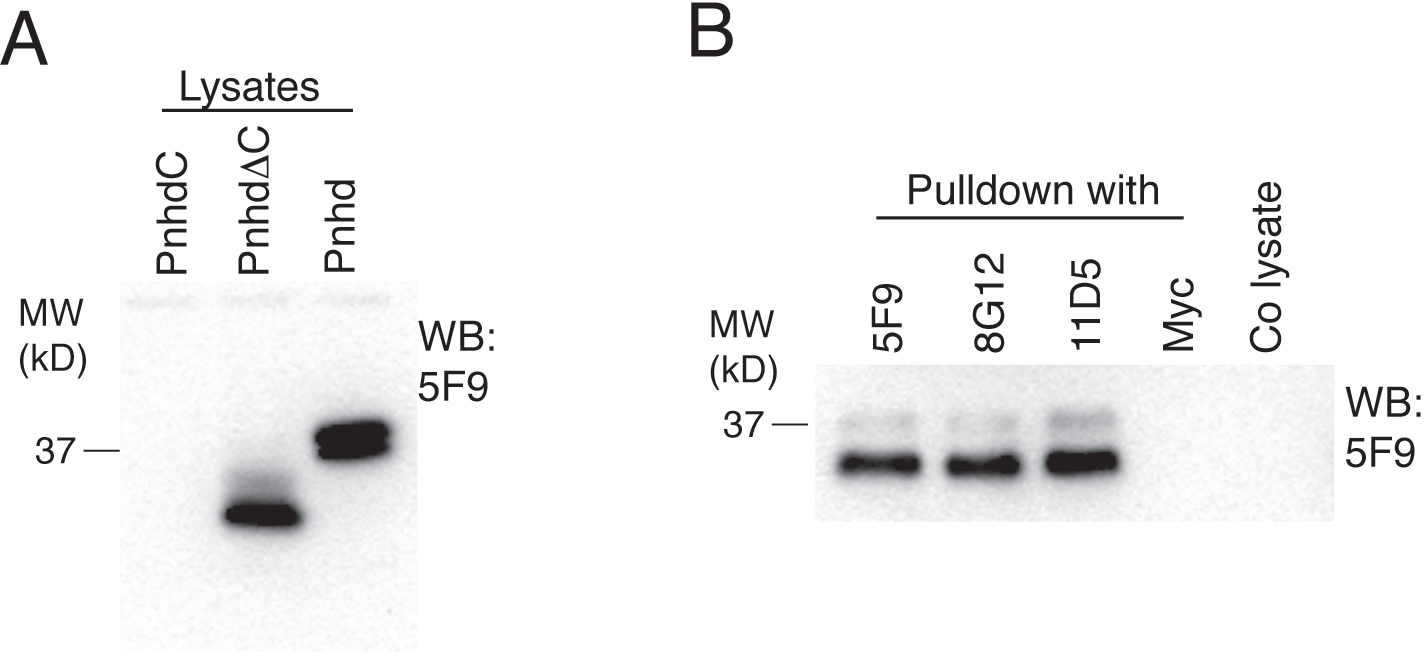
Both exogenous and endogenous Pnhd are recognized by specific antibodies, related to Figure 1. A, Embryos were injected with *flag-pnhd* RNA (75 pg) and the lysates of the injected and control uninjected embryos at stage 11.5-12 were immunoblotted with 5F9 antibodies. There is no visible signal in control embryos, but a strong 35-37 kDa band is detected in pnhd-expressing embryos. B, Lysates were prepared from normal embryos at stage 12, immunoprecipitated with the Pnhd-specific antibodies 5F9, 8G12,11D5 or anti-Myc as indicated and immunoblotted with 5F9.

**Suppl. Fig. 2.**
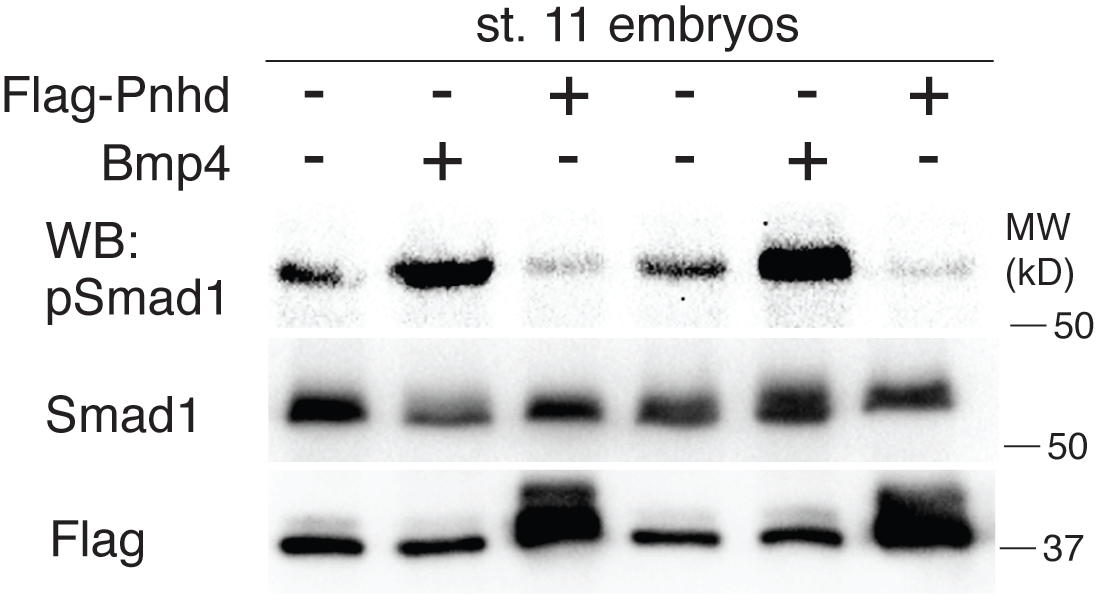
Inhibition of endogenous phospho-Smad1 levels by Pnhd, related to Figure 2. Two-cell embryos were injected into each blastomere with *flag-pnhd* RNA (1 ng) or *bmp4* RNA (50 pg). The injected embryos were collected when control siblings reached stage 11 for immunoblotting with anti-pSmad1, anti-Smad1 and anti-Flag antibodies. Pnhd reduced Smad1 phosphorylation whereas Bmp4 upregulated it.

**Suppl. Fig. 3.**
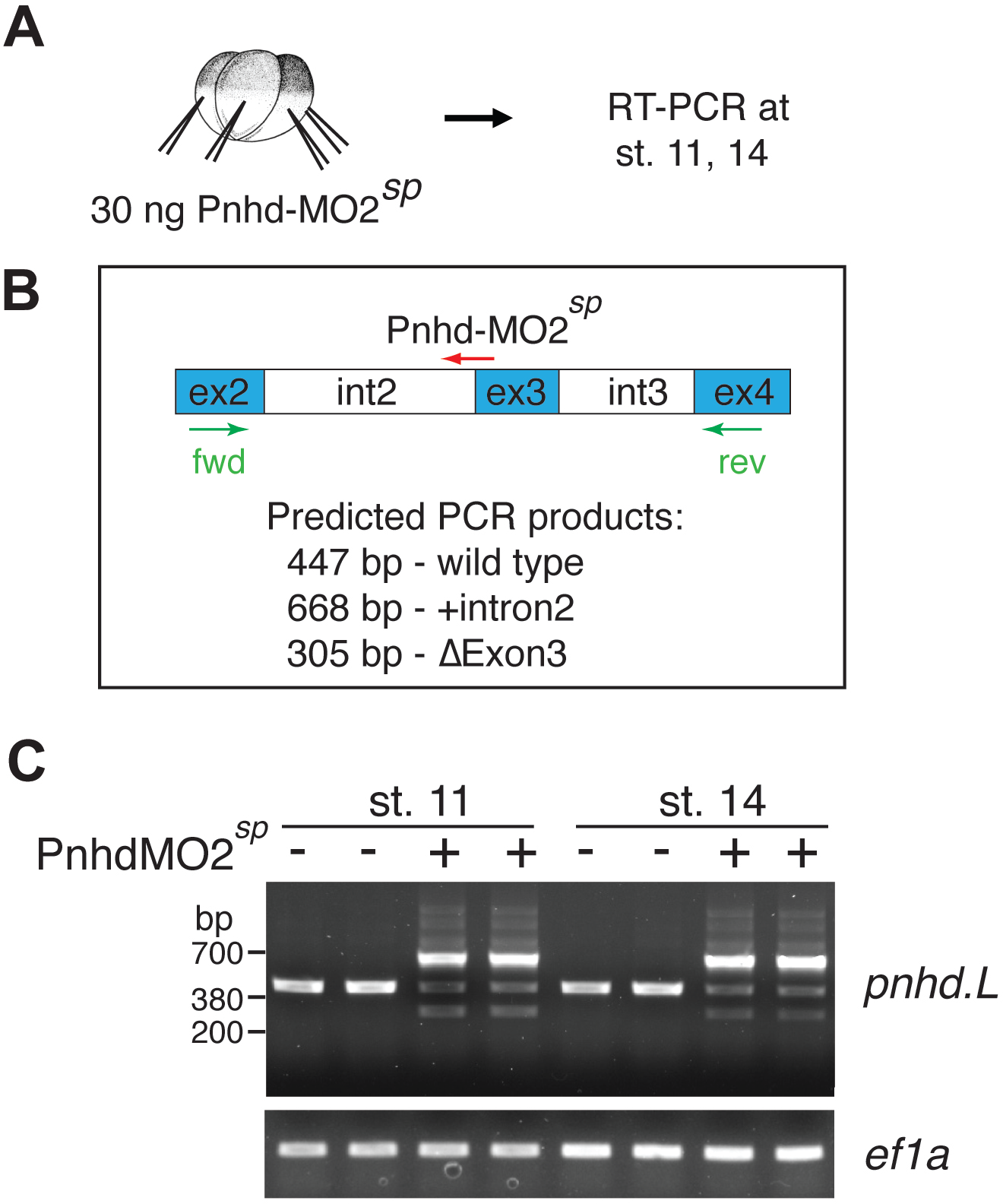
Characterization of the effects of PnhdMO2^sp^ on *pnhd* RNA splicing, related to Figure 4. A, Experimental scheme. Four-cell embryos were injected four times into marginal zone with 30 ng of PnhdMO2^sp^. RNAs were collected at stage 11 or 14 for RT-PCR. B, Pnhd-L gene structure with indicated exon (ex) and intron (int) boundaries. PnhdMO2^sp^ targets the junction of int2 and ex3. C, RT-PCR results show the addition of the intron 2 in the morphants. *ef1a* is a loading control.

**Suppl. Fig. 4.**
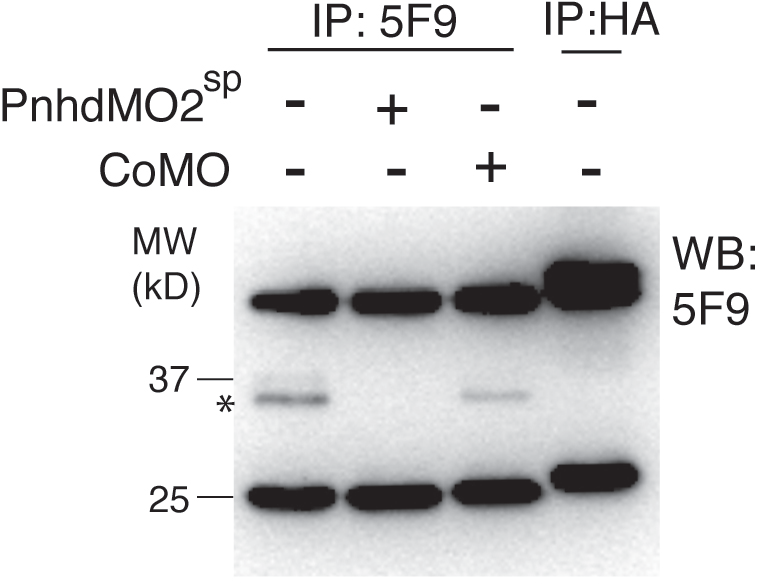
Validation of Pnhd knockdown by immunoprecipitation, related to Figure 4. Four-cell embryos were injected in the marginal zone with PnhdMO2^sp^ or control MO (40 ng each) as indicated. Sixty injected embryos were dissociated at stage 10 and cultured for 3 hrs. Conditioned media were immunoprecipitated with anti-Pnhd (5F9) or anti-HA antibodies and immunoblotted with 5F9. Pnhd-specific band has been eliminated in embryos injected with PnhdMO2^sp^.

**Suppl. Fig. 5.**
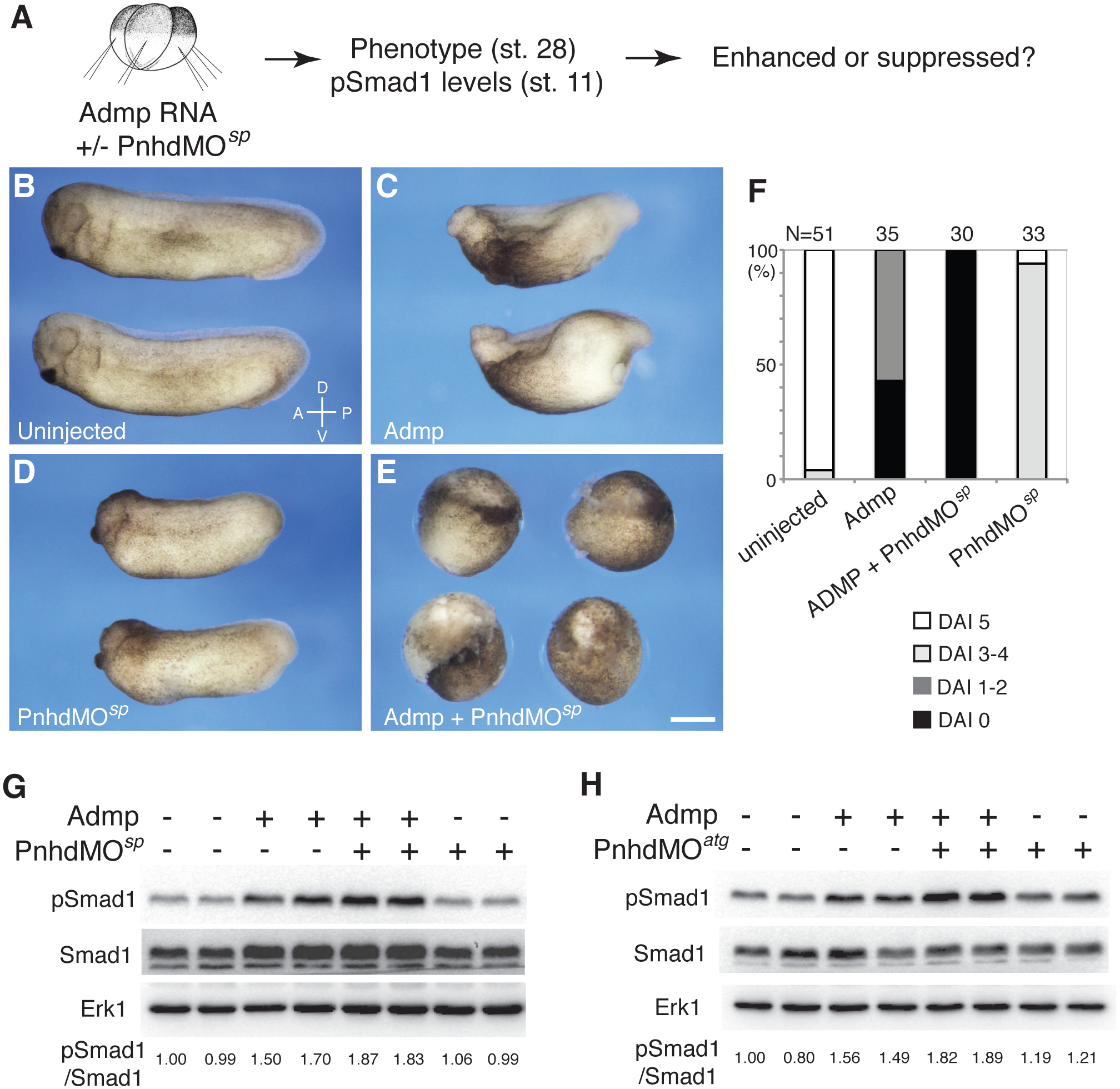
Enhancement of Admp ventralizing activity by PnhdMO^sp^, related to Figure 4. A, Experimental scheme. Marginal zone of four-cell embryos was injected with *admp* RNA (50 pg) or Pnhd MO^sp^ (60 ng) as indicated. B-E, PnhdMO^sp^ enhanced ventralization caused by Admp. Representative embryos are shown. F. Quantification of the data shown in B-E. Phenotypes were scored at stage 28 using dorsoanterior index (DAI). G, H, Immunoblotting of lysates of stage 11 embryos with anti-pSmad1, anti-Smad1 and anti-Erk1 antibodies. Normalized ratio of band intensities of pSmad1 relative to total Smad1 is indicated. Both PnhdMO^sp^ (60 ng) and PnhdMO^atg^ (10 ng) enhanced pSmad1 levels by Admp. Erk1 serves as a loading control. Data represent two to three independent experiments.

**Suppl. Fig. 6.**
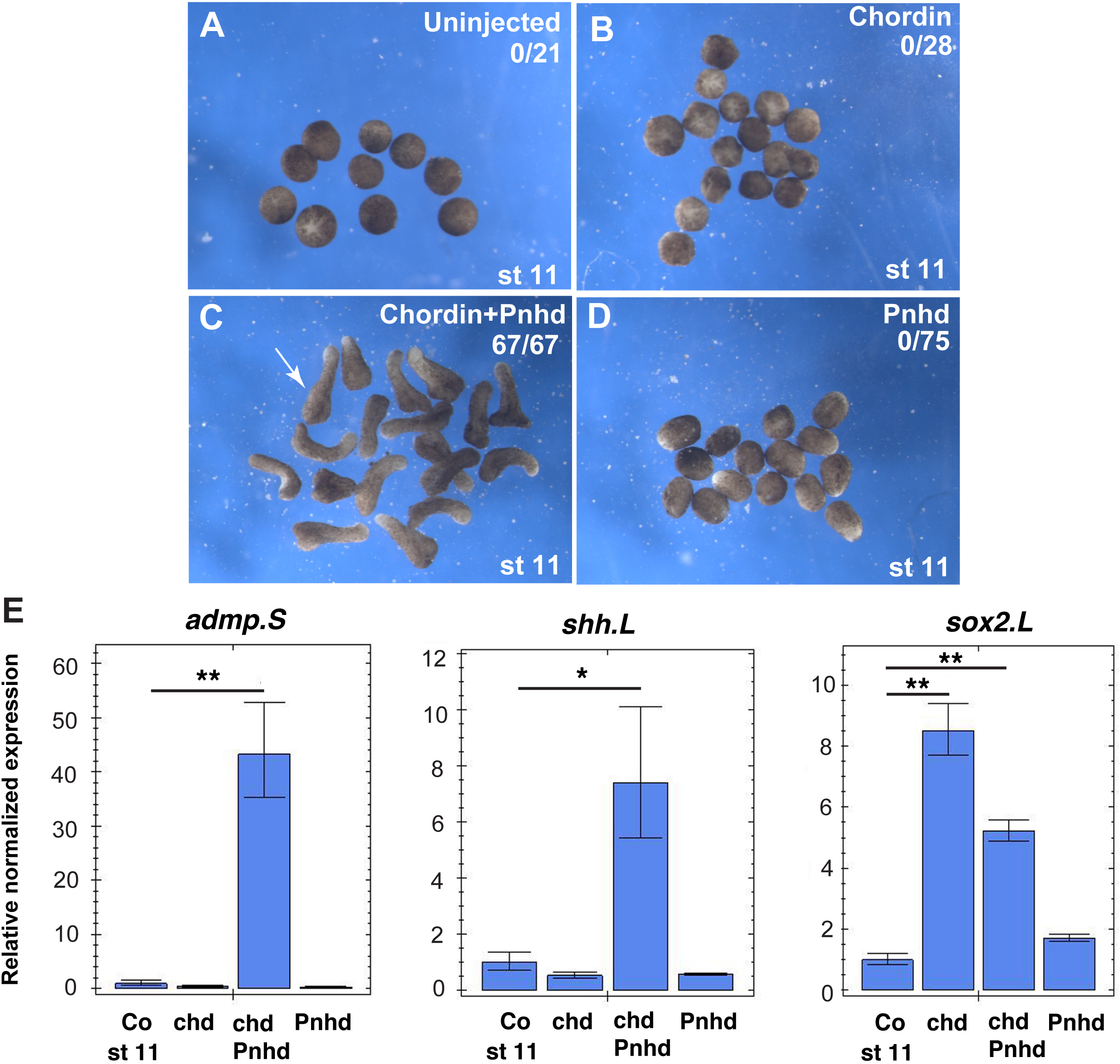
Pnhd promotes elongation of ectodermal explants in the presence of Chordin at midgastrula stage, related to Figure 5. A-D. Two cell embryos were injected with 0.5 ng of *pnhd* and/or 0.15 ng of *chordin* RNA as indicated. Ectodermal explants were dissected at stage 9.5-10 as shown in Fig. 5A. The explants were cultured until stage 11. E. Embryos were injected as described in A-D. Ectoderm explants were dissected at stages 9.5-10 and cultured until stage 11 to examine *admp* and *shh* by RT-qPCR. Data are representative of three experiments. Means +/- standard errors are shown. Significance was determined by the Student’s t-test, p<0.05 (*), p<0.01 (**).

## Notes

### Competing Interest Statement

The authors have declared no competing interest.

## References

Anders, S., and Huber, W. (2010). Differential expression analysis for sequence count data. Genome Biol 11, R106.

Bayramov, A.V., Eroshkin, F.M., Martynova, N.Y., Ermakova, G.V., Solovieva, E.A., and Zaraisky, A.G. (2011). Novel functions of Noggin proteins: inhibition of Activin/Nodal and Wnt signaling. Development 138, 5345-5356.

Bell, E., Munoz-­-Sanjuan, I., Altmann, C.R., Vonica, A., and Brivanlou, A.H. (2003). Cell fate specification and competence by Coco, a maternal BMP, TGFbeta and Wnt inhibitor. Development 130, 1381-1389.

Ben-­-Zvi, D., Shilo, B.Z., Fainsod, A., and Barkai, N. (2008). Scaling of the BMP activation gradient in Xenopus embryos. Nature 453, 1205-1211.

Cheng, A.M., Thisse, B., Thisse, C., and Wright, C.V. (2000). The lefty-­-related factor Xatv acts as a feedback inhibitor of nodal signaling in mesoderm induction and L-­-R axis development in xenopus. Development 127, 1049-1061.

Corallo, D., Schiavinato, A., Trapani, V., Moro, E., Argenton, F., and Bonaldo, P. (2013). Emilin3 is required for notochord sheath integrity and interacts with Scube2 to regulate notochord-­-derived Hedgehog signals. Development 140, 4594-4601.

Dale, L., Howes, G., Price, B.M., and Smith, J.C. (1992). Bone morphogenetic protein 4: a ventralizing factor in early Xenopus development. Development 115, 573-585.

De Robertis, E.M., and Kuroda, H. (2004). Dorsal-­-ventral patterning and neural induction in Xenopus embryos. Annu Rev Cell Dev Biol 20, 285-308.

Dollar, G.L., Weber, U., Mlodzik, M., and Sokol, S.Y. (2005). Regulation of Lethal giant larvae by Dishevelled. Nature 437, 1376-1380.

Dosch, R., and Niehrs, C. (2000). Requirement for anti-­-dorsalizing morphogenetic protein in organizer patterning. Mech Dev 90, 195-203.

Eroshkin, F.M., Nesterenko, A.M., Borodulin, A.V., Martynova, N.Y., Ermakova, G.V., Gyoeva, F.K., Orlov, E.E., Belogurov, A.A., Lukyanov, K.A., Bayramov, A.V., et al. (2016). Noggin4 is a long-­-range inhibitor of Wnt8 signalling that regulates head development in Xenopus laevis. Sci Rep 6, 23049.

Faure, S., Lee, M.A., Keller, T., ten Dijke, P., and Whitman, M. (2000). Endogenous patterns of TGFbeta superfamily signaling during early Xenopus development. Development 127, 2917-2931.

Graff, J.M., Thies, R.S., Song, J.J., Celeste, A.J., and Melton, D.A. (1994). Studies with a Xenopus BMP receptor suggest that ventral mesoderm-­-inducing signals override dorsal signals in vivo. Cell 79, 169-179.

Harland, R., and Gerhart, J. (1997). Formation and function of Spemann’s organizer. Annu Rev Cell Dev Biol 13, 611-667.

Harland, R.M. (1991). *In situ* hybridization: an improved whole-­-mount method for *Xenopus* embryos. In Methods Cell Biol, B.K. Kay, and H.B. Peng, eds. (San Diego: Academic Press Inc.), pp. 685-­-695.

Hopwood, N.D., Pluck, A., and Gurdon, J.B. (1989). MyoD expression in the forming somites is an early response to mesoderm induction in Xenopus embryos. EMBO J 8, 3409-3417.

Hsu, D.R., Economides, A.N., Wang, X., Eimon, P.M., and Harland, R.M. (1998). The Xenopus dorsalizing factor Gremlin identifies a novel family of secreted proteins that antagonize BMP activities. Mol Cell 1, 673-683.

Imai, K.S., Daido, Y., Kusakabe, T.G., and Satou, Y. (2012). Cis-­-acting transcriptional repression establishes a sharp boundary in chordate embryos. Science 337, 964-967.

Inomata, H., Shibata, T., Haraguchi, T., and Sasai, Y. (2013). Scaling of dorsal-­-ventral patterning by embryo size-­-dependent degradation of Spemann’s organizer signals. Cell 153, 1296-1311.

Inui, M., Montagner, M., Ben-­-Zvi, D., Martello, G., Soligo, S., Manfrin, A., Aragona, M., Enzo, E., Zacchigna, L., Zanconato, F., et al. (2012). Self-­-regulation of the head-­-inducing properties of the Spemann organizer. Proc Natl Acad Sci U S A 109, 15354-15359.

Itoh, K., Brott, B.K., Bae, G.U., Ratcliffe, M.J., and Sokol, S.Y. (2005). Nuclear localization is required for Dishevelled function in Wnt/beta-­-catenin signaling. J Biol 4, 3.

Joubin, K., and Stern, C.D. (1999). Molecular interactions continuously define the organizer during the cell movements of gastrulation. Cell 98, 559-571.

Kao, K.R., and Elinson, R.P. (1988). The entire mesodermal mantle behaves as Spemann’s organizer in dorsoanterior enhanced Xenopus laevis embryos. Dev Biol 127, 64-77.

Kenwrick, S., Amaya, E., and Papalopulu, N. (2004). Pilot morpholino screen in Xenopus tropicalis identifies a novel gene involved in head development. Dev Dyn 229, 289-299.

Kiecker, C., Bates, T., and Bell, E. (2016). Molecular specification of germ layers in vertebrate embryos. Cell Mol Life Sci 73, 923-947.

Kim, D., Langmead, B., and Salzberg, S.L. (2015). HISAT: a fast spliced aligner with low memory requirements. Nat Methods 12, 357-360.

Kimelman, D. (2006). Mesoderm induction: from caps to chips. Nat Rev Genet 7, 360-372.

Kjolby, R.A.S., and Harland, R.M. (2017). Genome-­-wide identification of Wnt/beta-­-catenin transcriptional targets during Xenopus gastrulation. Dev Biol 426, 165-175.

Kumano, G., Ezal, C., and Smith, W.C. (2006). ADMP2 is essential for primitive blood and heart development in Xenopus. Dev Biol 299, 411-­-423.

Lee, H.X., Ambrosio, A.L., Reversade, B., and De Robertis, E.M. (2006). Embryonic dorsal-­-ventral signaling: secreted frizzled-­-related proteins as inhibitors of tolloid proteinases. Cell 124, 147-159.

Marques, S., Borges, A.C., Silva, A.C., Freitas, S., Cordenonsi, M., and Belo, J.A. (2004). The activity of the Nodal antagonist Cerl-­-2 in the mouse node is required for correct L/R body axis. Genes Dev 18, 2342-2347.

Mayor, R., Morgan, R., and Sargent, M.G. (1995). Induction of the prospective neural crest of Xenopus. Development 121, 767-777.

Meno, C., Shimono, A., Saijoh, Y., Yashiro, K., Mochida, K., Ohishi, S., Noji, S., Kondoh, H., and Hamada, H. (1998). lefty-­-1 is required for left-­-right determination as a regulator of lefty-­-2 and nodal. Cell 94, 287-297.

Montague, T.G., Gagnon, J.A., and Schier, A.F. (2018). Conserved regulation of Nodal-­-mediated left-­-right patterning in zebrafish and mouse. Development 145.

Moos, M., Jr., Wang, S., and Krinks, M. (1995). Anti-­-dorsalizing morphogenetic protein is a novel TGF-­-beta homolog expressed in the Spemann organizer. Development 121, 4293-4301.

Muraoka, O., Shimizu, T., Yabe, T., Nojima, H., Bae, Y.K., Hashimoto, H., and Hibi, M. (2006). Sizzled controls dorso-­-ventral polarity by repressing cleavage of the Chordin protein. Nat Cell Biol 8, 329-338.

Niehrs, C. (2004). Regionally specific induction by the Spemann-­-Mangold organizer. Nat Rev Genet 5, 425-434.

Nieuwkoop, P.D., and Faber, J. (1967). Normal Table of Xenopus laevis (Daudin) (Amsterdam: North Holland).

Ossipova, O., Itoh, K., Radu, A., Ezan, J., and Sokol, S.Y. (2020). Pinhead signaling regulates mesoderm heterogeneity via the FGF receptor-­-dependent pathway. Development 147.

Peng, H.B. (1991). Xenopus laevis: Practical uses in cell and molecular biology. Solutions and protocols. Methods Cell Biol 36, 657-662.

Peyrot, S.M., Wallingford, J.B., and Harland, R.M. (2011). A revised model of Xenopus dorsal midline development: differential and separable requirements for Notch and Shh signaling. Dev Biol 352, 254-266.

Piccolo, S., Agius, E., Leyns, L., Bhattacharyya, S., Grunz, H., Bouwmeester, T., and De Robertis, E.M. (1999). The head inducer Cerberus is a multifunctional antagonist of Nodal, BMP and Wnt signals. Nature 397, 707-710.

Piccolo, S., Sasai, Y., Lu, B., and De Robertis, E.M. (1996). Dorsoventral patterning in Xenopus: inhibition of ventral signals by direct binding of chordin to BMP-­-4. Cell 86, 589-598.

Reversade, B., and De Robertis, E.M. (2005). Regulation of ADMP and BMP2/4/7 at opposite embryonic poles generates a self-­-regulating morphogenetic field. Cell 123, 1147-1160.

Sasai, Y., Lu, B., Steinbeisser, H., and De Robertis, E.M. (1995). Regulation of neural induction by the Chd and Bmp-­-4 antagonistic patterning signals in Xenopus. Nature 376, 333-336.

Sasai, Y., Lu, B., Steinbeisser, H., Geissert, D., Gont, L.K., and De Robertis, E.M. (1994). Xenopus chordin: a novel dorsalizing factor activated by organizer-­-specific homeobox genes. Cell 79, 779-790.

Schohl, A., and Fagotto, F. (2002). Beta-­-catenin, MAPK and Smad signaling during early Xenopus development. Development 129, 37-52.

Suzuki, A., Thies, R.S., Yamaji, N., Song, J.J., Wozney, J.M., Murakami, K., and Ueno, N. (1994). A truncated bone morphogenetic protein receptor affects dorsal-­-ventral patterning in the early Xenopus embryo. Proc Natl Acad Sci U S A 91, 10255-10259.

Tuazon, F.B., and Mullins, M.C. (2015). Temporally coordinated signals progressively pattern the anteroposterior and dorsoventral body axes. Semin Cell Dev Biol 42, 118-­-133.

von Dassow, G., Schmidt, J.E., and Kimelman, D. (1993). Induction of the Xenopus organizer: expression and regulation of Xnot, a novel FGF and activin-­-regulated homeo box gene. Genes Dev 7, 355-366.

Willot, V., Mathieu, J., Lu, Y., Schmid, B., Sidi, S., Yan, Y.L., Postlethwait, J.H., Mullins, M., Rosa, F., and Peyrieras, N. (2002). Cooperative action of ADMP-­- and BMP-­-mediated pathways in regulating cell fates in the zebrafish gastrula. Dev Biol 241, 59-78.

Yan, Y., Ning, G., Li, L., Liu, J., Yang, S., Cao, Y., and Wang, Q. (2019). The BMP ligand Pinhead together with Admp supports the robustness of embryonic patterning. Sci Adv 5, eaau6455.

